# A Pumpless Microfluidic Neonatal Lung Assist Device for Support of Preterm Neonates in Respiratory Distress

**DOI:** 10.1101/2020.02.22.961102

**Authors:** Mohammadhossein Dabaghi, Niels Rochow, Neda Saraei, Gerhard Fusch, Shelley Monkman, Kevin Da, Alireza Shahin-Shamsabadi, John L. Brash, Dragos Predescu, Kathleen Delaney, Christoph Fusch, P. Ravi Selvaganapathy

## Abstract

Premature neonates suffer from respiratory morbidity as their lungs are immature and current supportive treatment such as mechanical ventilation or extracorporeal membrane oxygenation (ECMO) cause iatrogenic injuries. A non-invasive and biomimetic concept known as the “artificial placenta” would be beneficial to overcome complications associated with the current respiratory support of preterm infants. Here, a pumpless oxygenator connected to the systemic circulation supports the lung function to relieve respiratory distress. In this paper, we demonstrate the first successful operation of a microfluidic, artificial placenta type neonatal lung assist device (LAD) on a newborn piglet model which is the closest representation of preterm human infants. This LAD has high oxygenation capability in both pure oxygen and room air as the sweep gas. It was able to relieve the respiratory distress that the newborn piglet was put under during experimentation, repeatedly and over significant duration of time. These findings indicate that this LAD has potential application as a biomimetic artificial placenta to support respiratory needs of preterm neonates.

## Introduction

Premature births are associated with significant respiratory morbidity and mortality. Neonates, especially those with very low birthweights and born at less than 28 weeks of gestational age^[1,2]^ have a very low survival rate. In particular, those born earlier than 24 weeks of gestational age^[3]^ have more than 50 % mortality. One of the leading causes of morbidity associated with preterm birth is respiratory failure caused by lack of surfactants in immature lungs as well lung hypoplasia. ^[4–7]^. Mechanical ventilation is often used for respiratory support of such premature neonates^[8]^ who are in respiratory failure. Nonetheless, mechanical ventilation is invasive and associated with severe complications such as pulmonary injury, chronic lung disease, and related diseases of prematurity such as retinopathy of prematurity, intraventricular hemorrhage or necrotizing enterocolitis which would lead to several long-term side effects^[9–11]^. In late-preterm and term infants, extracorporeal membrane oxygenation (ECMO) could be an alternative choice of treatment. However, ECMO is invasive and requires central vascular access and therefore, surgery. Further, the current hollow fiber based blood oxygenators that are used in ECMO are not suitable for neonatal application due to their high priming volume (> 20 mL), high pressure drop which requires an external pump for perfusion, and their gas exchange dependency on oxygen^[8,12]^. Moreover, other complications such as cerebral injury with intracranial hemorrhage or stroke, poor somatic growth, and poor neurodevelopment can be caused by this technology^[13]^.

Microfabrication technologies have been employed to fabricate blood oxygenators that can address some of the limitations of current ECMO devices by using biomimetic architecture similar to the vascular network in the lung in order to enhance gas transfer efficiency, increase effective surface area for transfer as well as reduce shear stress in blood flow and avoid the formation of blood stagnation zones^[14,15]^. Several microfluidic blood oxygenators^[12,16,25–32,17–24]^ have been introduced aimed at improving gas exchange efficiency but all of them use external pumps to perfuse the blood through them. An alternate approach is to use the arterio-venous pressure difference in the body as a natural pumping mechanism to circulate the blood through an oxygenator – a concept known as the artificial placenta. The artificial placenta (AP) approach is characterized by (i) blood circulating through a passive device pumped solely by the arteriovenous pressure difference of the neonate’s heart and (ii) being capable of exchanging blood gases with the room air. This concept is physiologically similar to in-utero conditions and overcomes some of the issues associated with traditional ECMO devices^[8,12,16]^. Traditionally used hollow fiber ECMO devices have high priming volume and are not suitable for neonatal application. Only some of microfluidic oxygenators^[12,16,29–31]^ have been explicitly designed for use as an artificial placenta device. Most artificial placenta-type microfluidic blood oxygenators have been only been examined in vitro^[12,29,30]^ and even those^[16,31]^ tested in vivo have not been shown to provide adequate oxygenation in room air to support a suitable animal model in respiratory distress.

Here, we present the first high-performance, pumpless, neonatal lung assist device that in *in-vivo* experiments on newborn piglets (of similar size to neonates) has demonstrated its ability to overcome respiratory distress. The LAD was assembled from high-performance microfluidic blood oxygenators (MBOs)^[30]^ in a parallel configuration to enable sufficient flow of blood through it as it was pumped by the pressure generated solely by the piglet’s heart. Additionally, the entire LAD was coated with heparin to reduce its systemic infusion and the efficacy of this coating was evaluated. We report the in-vitro gas exchange performance of the LAD as well as its in-vivo performance when the piglet was intentionally put in respiratory distress by placing it under hypoxic conditions. Systemic blood gas measurements as well as cardiovascular conditions were monitored during repeated cycles of hypoxic and normoxic conditions over a period of 200 minutes. Our results showed that this passive pumpless LAD could provide adequate oxygenation in room air to restore near normoxic condition and has the potential to meet the clinical requirements to support neonates under respiratory distress.

## Results

### In vitro testing of LAD using bovine blood

The gas exchange performance of the LAD in-vitro was investigated in enriched oxygen environment as well as in room air using heparinized (3 unit mL^-1^) bovine blood. The oxygen saturation levels (Sa O_2_) of the blood were adjusted to 58 ± 2 % for experiments performed in an enriched oxygen environment and 63 ± 1 % for experiments conducted in room air. The desaturated blood was pumped through the device and the oxygen saturation was measured in the outlet.

The LAD itself was assembled from several MBOs (16 in parallel) with blood channel height of 180 μm which were made of PDMS (methods section). The LAD was able to fully oxygenate blood (Sa O_2_ = ~ 100 %) up to a flow rate of 80 mL min^-1^ in an enriched oxygen environment and to increase Sa O_2_ to ~ 95 % at a blood flow rate of 100 mL min^-1^ (Figure 1a). The oxygen uptake was calculated by summing the amount of dissolved oxygen and adsorbed oxygen to red blood cells^[12]^. The oxygen uptake was between 0.92 to 5.71 mL min^-1^ with a linear increase with raising blood flow rates while the LAD was using oxygen as the sweep gas (Figure 1b). The LAD had a different gas transfer performance in room air as oxygen concentration was lower. Consequently, oxygen saturation level achieved was lower and the blood was fully oxygenated only up to a blood flow rate of 20 mL min^-1^. The outlet saturation level dropped to 87 % at a blood flow rate of 60 mL min^-1^(Figure 1c). The oxygen uptake linearly increased with blood flow rates and was 1.82 mL min^-1^ at a blood flow rate of 60 mL min^-1^ in room air (Figure 1d). The pressure drop of the LAD (Figure 1e), as well as the LAD with the extracorporeal circuit (Figure 1f), was measured over a range of flow rates from 10 – 100 mL min^-1^. Carbon dioxide release data is presented in the supplementary Figure S1.

**Figure 1:**
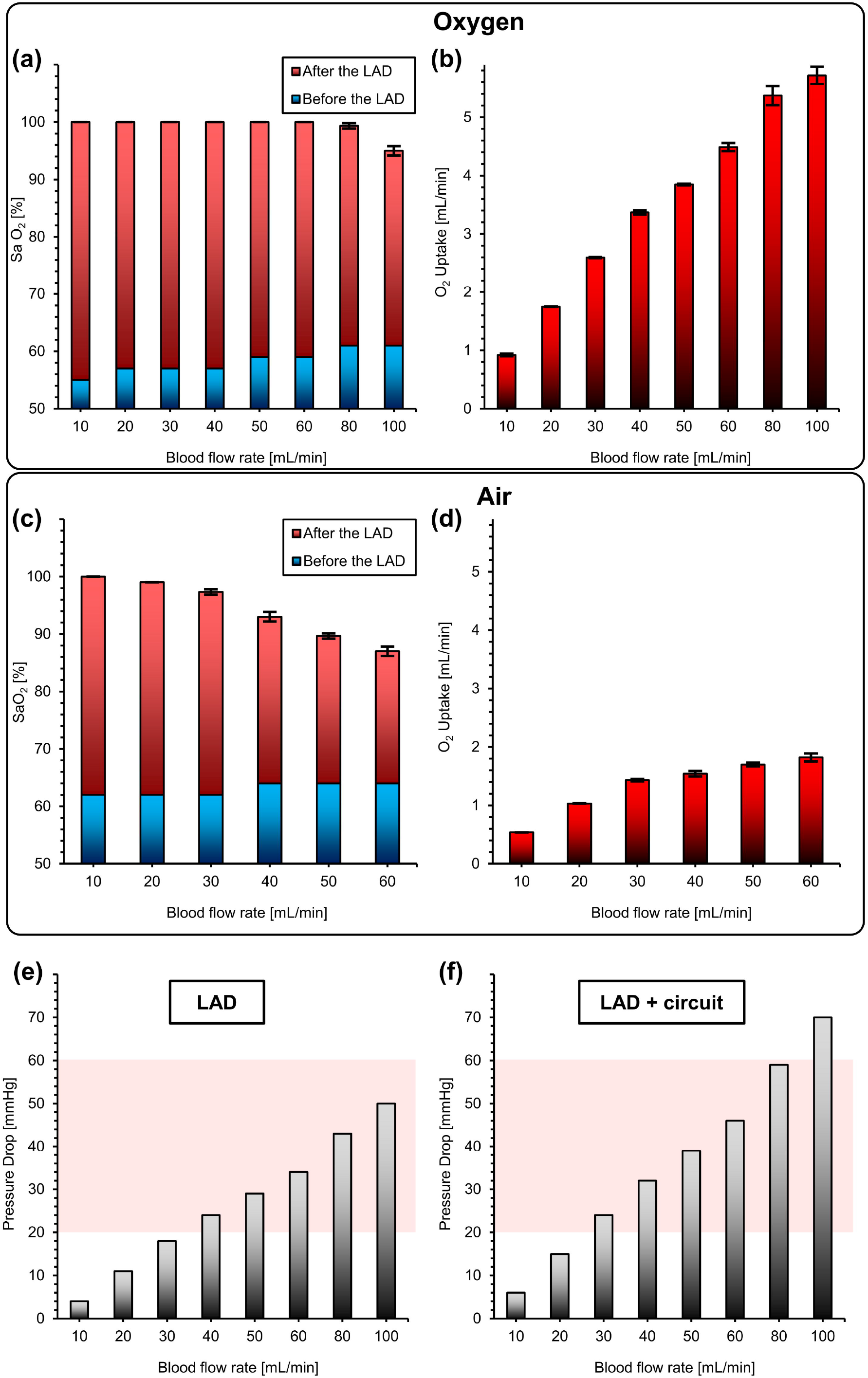
In vitro test of the LAD with heparinized bovine blood at various blood flow rates: (a) oxygen saturation level before and after the LAD in oxygen, (b) oxygen uptake in oxygen (c) oxygen saturation level before and after the LAD in room air, (d) oxygen uptake in room air, (e) pressure drops for the LAD (the shaded region indicates the operational pressure drops range for a pumpless operation), and f) the pressure drops for the LAD with the extracorporeal circuit. Data are means ± SD, n = 3. Hematocrit was 38 % and 29 % for oxygen and air conditions, respectively.

We also tested LAD that had MBOs with the blood channel’s height of 130 μm and compared it with the current (180 μm) design. This LAD was tested in vitro and in vivo using only pure oxygen as the sweep gas. Interestingly, both LADs (one with microchannel’s height of 130 μm and another one had a microchannel’s height of 180 μm) had similar oxygenation performance (Figure S9a in the supplementary) but noticeably different hydraulic resistance. The LAD with 130 μm channel height had lower blood flow rate (60-70 mL/min) in the corresponding in-vivo experiment (Figure S9b in the supplementary). Nevertheless, even this design was able to achieve nearly 90% systemic oxygenation levels using hypoxic conditions. Based on this comparison, the LAD with MBOs having a channel height of 180 μm was used in the subsequent long-term experiment as the optimized configuration (the details of the animal experiment for the LAD with the height of 130 μm are presented in the supplementary Figures S10 – S12).

### Modification of blood-in-contact surfaces with heparin

PDMS is hydrophobic and is known to adsorb proteins readily from blood^[33]^, leading to the activation of blood coagulation, platelet adhesion and thrombosis. To minimize these effects, all blood-contacting surfaces were coated with heparin using the set-up depicted in Figure 2a1. The coating process was optimized using fluorescently labeled heparin which allowed assessment of the quality and stability of the coating. For instance, the wash time to remove residual heparin was optimized to 15 min by monitoring the fluorescence intensity of the wash eluate (Figure 2a2).

**Figure 2:**
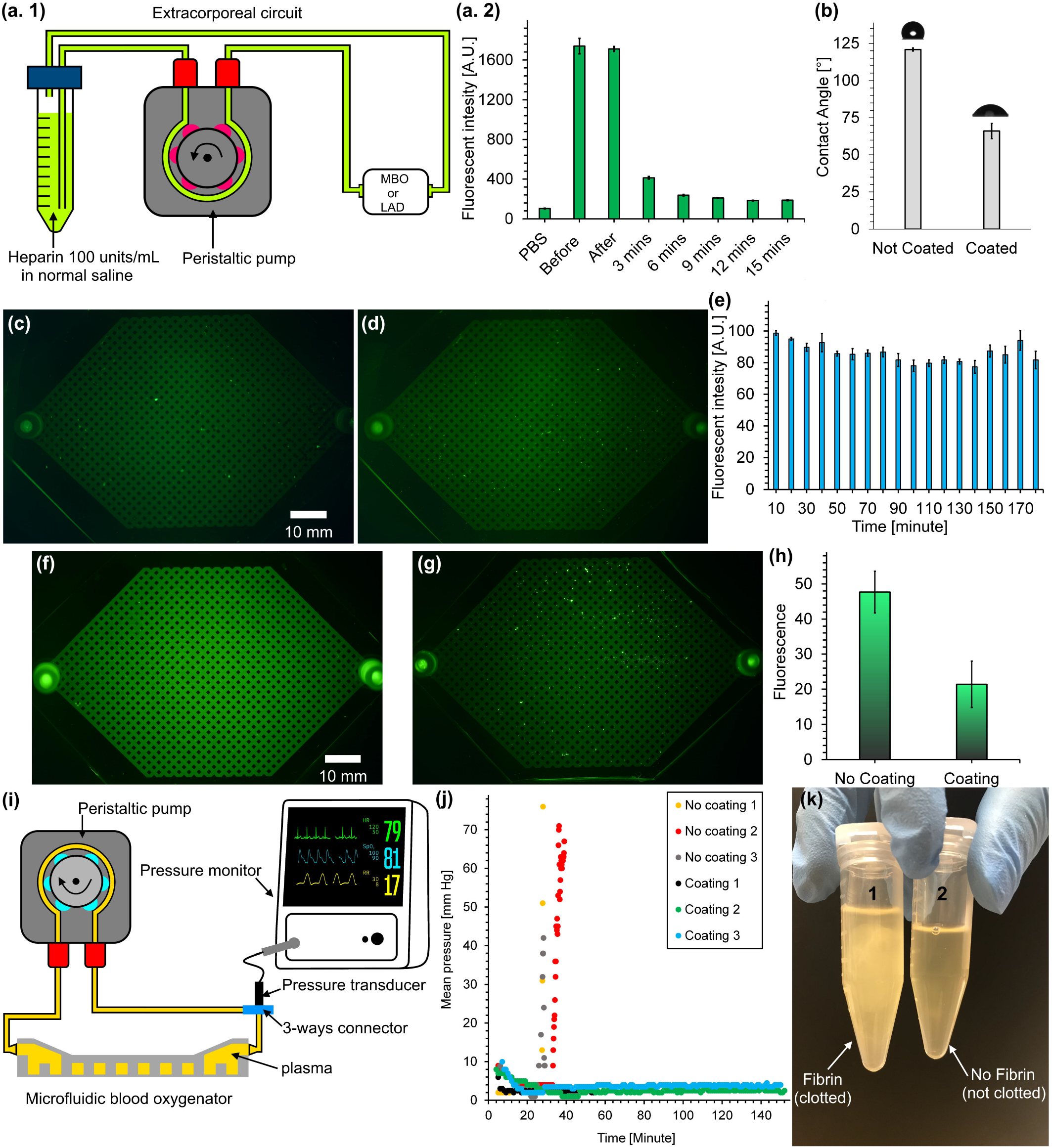
In vitro experiments to assess the effects of heparin coating on LADs or MBOs. (a. 1) experimental setup for coating, (a. 2) fluorescence intensity of solution samples from washing an FITC-heparin coated MBO with PBS, (b) water contact angles of coated and non-coated surfaces, (c) wide-angle image of an MBO coated with FITC-heparin, (d) wide-angle image of an MBO coated with FITC-heparin after contact with flowing PBS for 20 h, (e) fluorescence intensity of solution samples collected during contact with a heparin-coated MBO for 3 h, (f wide-angle image of FITC-fibrinogen adsorbed to an uncoated MBO, (g) wide-angle image of FITC-fibrinogen adsorbed to a heparin-coated MBO, (h) fluorescence intensities of uncoated and heparin-coated surfaces obtained using ImageJ software, (i) experimental setup for pressure measurement during plasma circulation, (j) representative pressure trace during plasma circulation in coated and uncoated MBOs (n = 3), and (k) collected plasma after circulation in the MBOs with coated and uncoated surfaces.

Coating PDMS with heparin decreased the water contact angle from 120° to 66°, rendering the surface moderately hydrophilic and possibly reducing the potential for clot formation on the surface (Figure 2b). More importantly, heparin as an anticoagulant on the surface is expected to inhibit clotting via inhibition of thrombin.

Wide-angle fluorescence images of the MBO surfaces after heparin coating showed that the coating on the blood vascular network surfaces was uniform (Figure 2c). Subsequently, PBS was perfused in the MBO at a flow rate of 3 mL min^-1^ for 20 h to investigate the stability of the heparin coating. As seen in Figure 2d, fluorescence image of FITC-labeled heparin coated inside the MBO showed the similar pattern to the coated MBO before perfusion by PBS without any significant reduction in its intensity. The bonding mechanism of heparin to the PDMS is unknown but could involve hydrogen bonding between hydroxyl groups of heparin and the oxygen atoms in the PDMS^[34]^ as well as physical adsorption, thus giving a coating which appears to be stable in PBS for at least several hours. Furthermore, the heparin content was determined in PBS samples at 10 min intervals during the first three hours. It appeared that a small heparin quantity was removed initially but no additional loss occurred over the subsequent three hours period (Figure 2e). These data suggest that the heparin coating was sufficiently stable under the conditions to be used with the oxygenator device.

A solution of FITC-fibrinogen and PBS buffer (2 mg mL^-1^) was circulated to coated and non-coated MBOs to visualize the rate of the protein adsorption on the non-coated and coated surfaces (Figure 2f and g). The fluorescence intensity of each MBO was quantified using ImageJ software which showed a significant difference in fibrinogen adsorption between coated and non-coated MBOs suggesting that the heparin coating was effective in reducing fibrinogen adsorption (Figure 2h).

### Heparin coated MBOs do not exhibit appreciable clot formation in plasma

The formation of micro-clots can impact device performance. This is especially true for a device with microchannels that could be easily occluded by micro-clots, resulting in uneven flow distribution, decreased gas transfer rates, and possible device failure^[35,36]^. To investigate the possible formation of micro-clots, a plasma clotting assay was performed on both heparin-coated and uncoated MBOs using recalcified citrated plasma (Figure 2i). The pressure in circuits containing uncoated MBOs increased sharply after 29 ± 3.5 minutes. (By comparison recalcified plasma stored in a falcon tube clotted after about 20 minutes.) In contrast, no perceptible change in pressure was observed in the loops with coated MBOs up to 150 min demonstrating the beneficial effect of the heparin coating (Figure 2j). These plasma clots were visible in the samples collected from the loops with uncoated devices, while the plasma in the loops with coated devices was clear (Figure 2k).

### Microfluidic LAD rescues hypoxic piglet from respiratory distress

The LAD was tested *in vivo* exposed to room air or to oxygen enriched air using sweep gas on a hypoxic piglet model. Piglets represent the human neonate more closely that any other animal model and was chosen for this experiment. The experimental setup was adapted from Rochow et. al^[16]^ (Figure 3) where the animal was connected arterio-venously to an extracorporeal circuit that has a short cut loop which returns the blood without any oxygenation and to the LAD in parallel. Initially, when the extracorporeal circuit was connected and perfused by the pressure difference provided by the piglet’s heart, the LAD loop stayed closed, while blood flowed through the shortcut line. Under normoxic condition this setting was used to ensure that all vital parameters were in steady state (termed Shortcut Normoxic). Hypoxic condition targeting peripheral oxygen saturation (SpO_2_) of 60% was simulated for the paralyzed piglet by adjusting the ventilator: PIP of 10 cm H_2_O, PEEP of 5 cm H_2_O, IT of 0.33 s, and RR of 15 – 19 breaths min^-1^. After achieving the target SpO_2_, immediately blood gases and cardiovascularly parameter were measured without LAD support while blood flowed through the shortcut (termed Shortcut Hypoxic). In order to bring the LAD online, the shortcut valve was closed and the LAD circuit valve opened redirecting the blood flow through the LAD loop (termed LAD Hypoxic). Error! Reference source not found. in the supplementary shows the sequence of conditions that the piglet was exposed to over the time frame for the in-vivo experiment. Initially, baseline measurements were made by exposing the piglet to normoxic condition with the blood flowing through the shortcut. Then, the hypoxic condition was initiated, and blood flow switched between the shortcut and through the LAD in three cycles each, when the LAD is exposed to pure oxygen and air. Multiple blood samples were taken under each of these conditions and at various time points as shown in Error! Reference source not found. in the supplementary. The blood samples were collected from the femoral artery, right atrium, before the LAD, and after the LAD to conduct blood gas analysis. At the end of the experiment, shortcut normoxic condition was established (control) and a final set of blood samples were taken for measurement of blood gas levels to ensure that the piglet recovered to the initial baseline and was not affected by the intervention.

**Figure 3:**
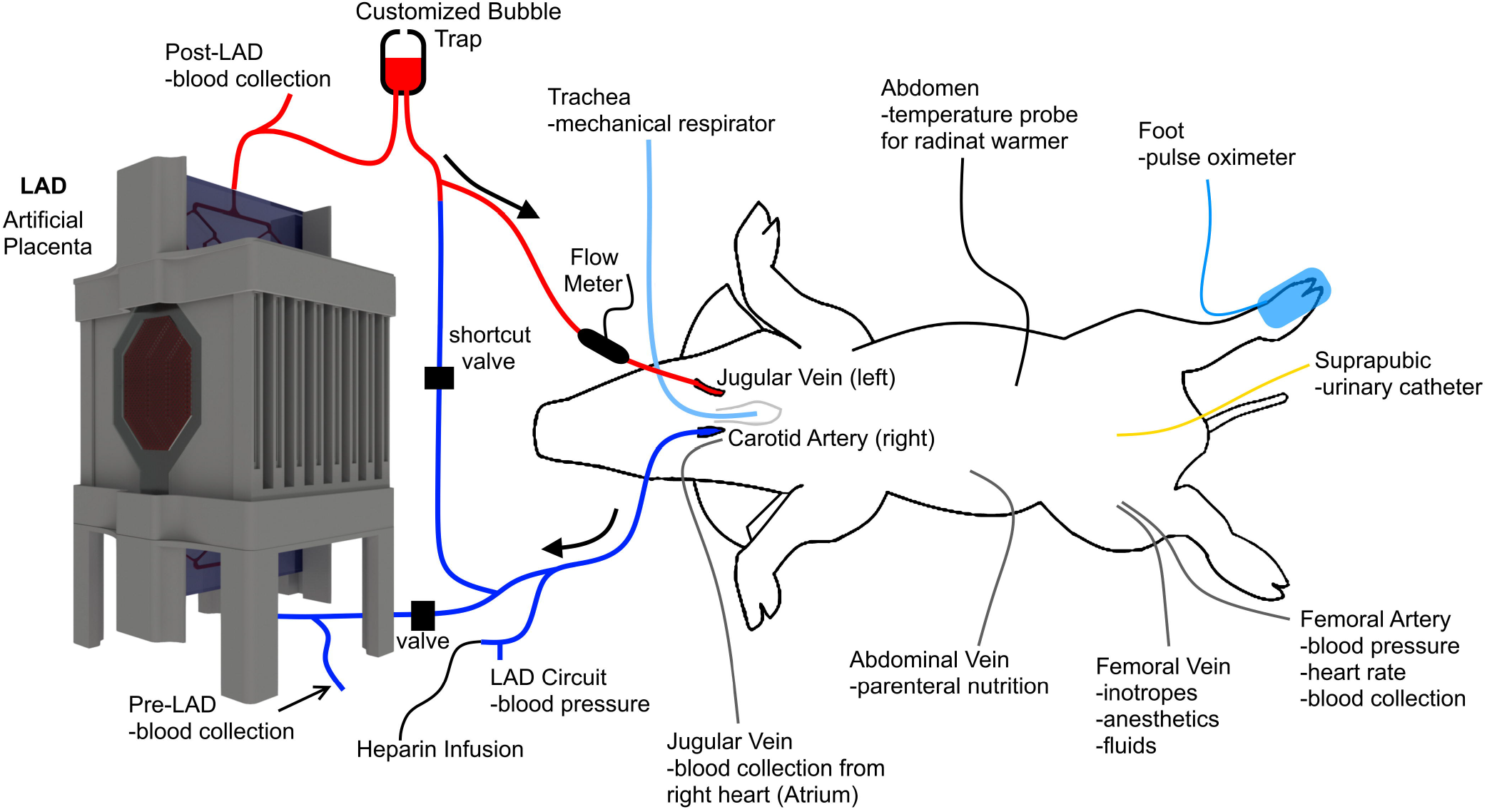
Experimental setup of the newborn piglet model for testing a pumpless neonatal lung assist device for artificial placenta application.

### Systematic blood measurements

It is vital to discern the effect of the extracorporeal circuit on systematic blood parameters such as blood flow rate, heart rate, mean LAD/shortcut pressure, and mean blood pressure. Since the LAD was connected to the systemic circulation (an arteriovenous connection) in a parallel configuration, the resistance to blood flow would be reduced which would result in a higher systemic blood flow rate (under short cut normoxic condition). When the piglet is made hypoxic, the systemic blood flow rate was found to increase as it compensates for the lower oxygen saturation level in blood (Figure 4a and e). However, when the extracorporeal blood circulation is switched from the shortcut to the LAD, it adds not only oxygen to the blood but also increases the resistance in the parallel extracorporeal circuit. This results in the observed lowering of the systemic blood flow rate when the LAD is connected. Heart rate was found to be elevated under the hypoxic condition to pump blood faster and increase the rate of gas exchange, but it dropped back to normal levels when the LAD became active indicating that the LAD was successful in alleviating the hypoxic condition (Figure 4b and f). The mean blood pressure at the location where the blood emerges extracorporeally (Figure 4c and g) was always lower in the shortcut hypoxic condition compared to the LAD hypoxic (the condition that had the LAD in the circuit) confirming that the LAD line achieved lower flow rates. Moreover, mean systemic blood pressure (at the femoral artery) (Figure 4d and h) was elevated under the hypoxic conditions and returns to normal levels when oxygenation through the LAD is initiated.

**Figure 4:**
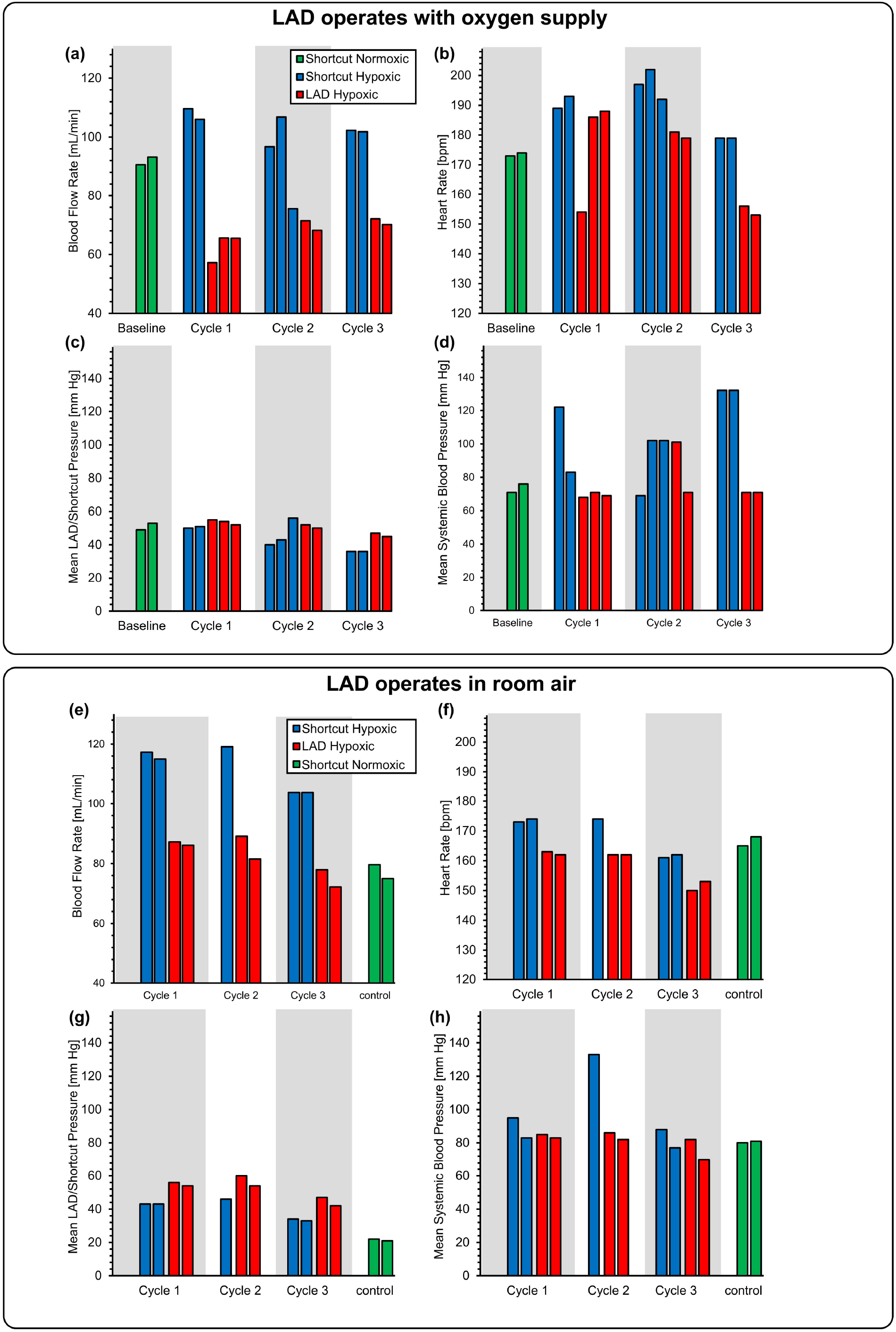
Effect of extracorporeal bypass on cardiovascular parameters: (a) blood flow rate while the LAD was using oxygen as the sweep gas, (b) heart rate while the LAD was using oxygen as the sweep gas, (c) mean LAD/shortcut pressure while the LAD was using oxygen as the sweep gas, (d) mean systemic arterial blood pressure measured at the femoral artery while the LAD was using oxygen as the sweep gas, (e) blood flow rate when the LAD was exposed to room air, (f) heart rate when the LAD was exposed to room air, (g) mean LAD/shortcut pressure when the LAD was exposed to room air, and (h) mean systemic arterial blood pressure measured at the femoral artery when the LAD was exposed to room air.

### In vivo gas exchange testing

At the end of each cycle, blood samples were collected extracorporeally before and after the LAD as well as from the femoral artery and the jugular vein from the right cardiac atrium to determine systemic blood gas levels. The saturation of oxygen (SaO_2_), partial pressure of oxygen (pO_2_), and partial pressure of carbon dioxide (pCO_2_) were measured in these samples by a blood gas analyzer. Under the shortcut hypoxic condition, the blood flow rate was stable and ranged from 102 mL min^-1^ to 110 mL min^-1^. When the LAD (LAD hypoxic condition) was connected, its hydraulic resistance was low enough for it to be perfused solely by the piglet’s heart and generate a stable blood flow rate from 65 mL min^-1^ to 72 mL min^-1^ (37 – 41 mL min^-1^ kg^-1^), clearly demonstrating that the design was able to support clinically relevant flow rates in a pumpless manner. When the LAD was placed in an enriched oxygen environment (a chamber with slight oxygen flow of ~ 15 L hr^-1^), the LAD was able to fully oxygenate the blood at these relevant flow rates in all experiments. In one instance, it was able to increase the Sa O_2_ from a low 78 % to 100% at a high blood flow rate of ~ 70 mL min^-1^ (Figure 5a). The LAD also could significantly increase the pO_2_ up to 473 mm Hg (as compared to 68 mm Hg at its inlet) resulting in highly oxygenated blood leaving the LAD outlet (Figure 5b). pCO_2_ was also decreased at this condition with a maximum drop of 81 mm Hg to 50 mm Hg at a blood flow rate of 70 mL min^-1^ (Figure 5c).

**Figure 5:**
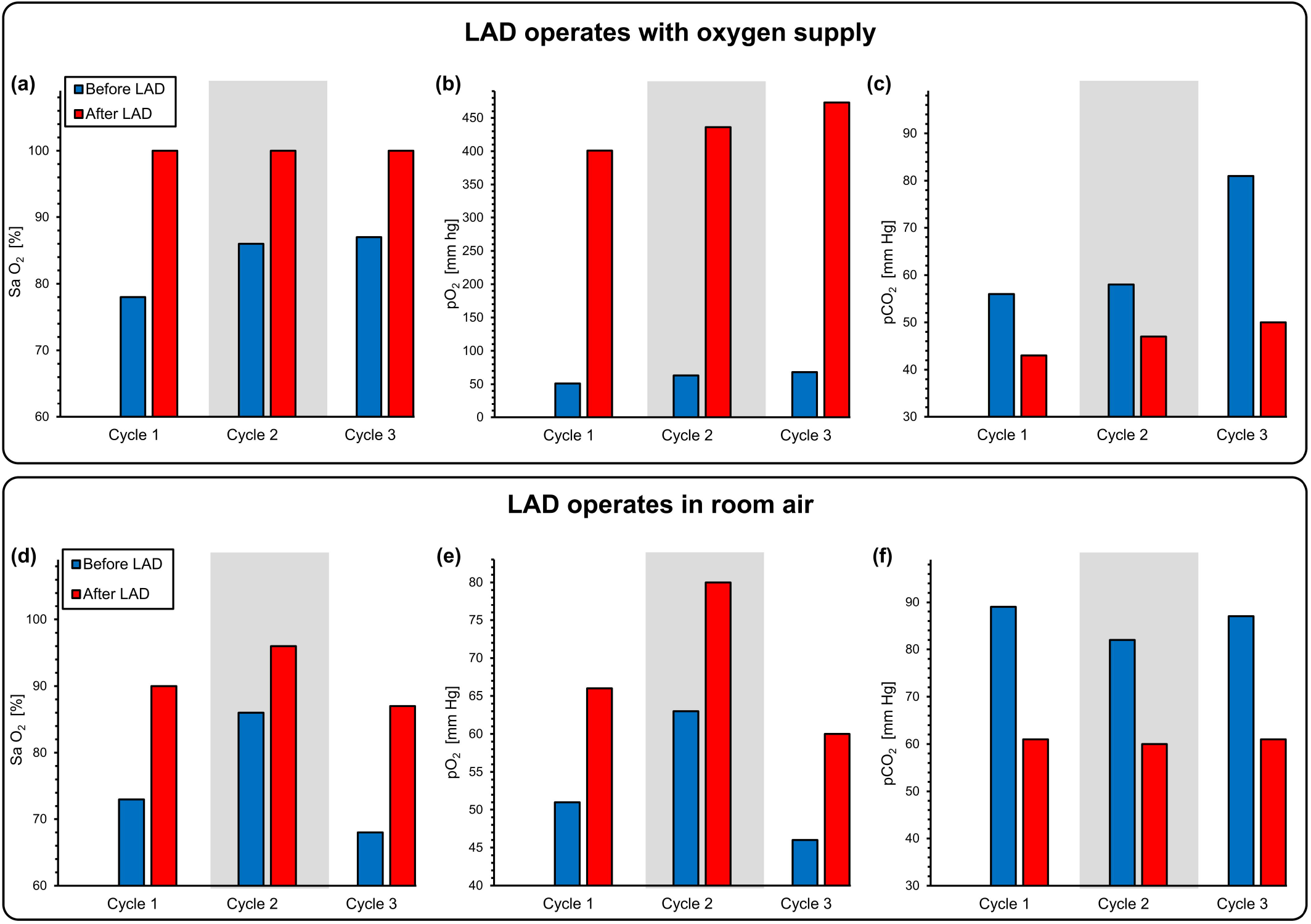
Gas exchange at the LAD when connected in-vivo to a piglet. Measurements were of the blood at the inlet and outlet of the LAD when it is connected to the piglet and pumped by the arterio-venous pressure difference: (a) Sa O_2_, (b) pO_2_, (c) pCO_2_ before and after the LAD using oxygen as the sweep gas, and (d) Sa O_2_, (e) pO_2_, f) pCO_2_ before and after the LAD exposed directly to room air.

When the LAD was exposed to ambient air, the gas exchange performance was lower but still significant. For instance, in one cycle, the LAD was capable of increasing the SaO_2_ from 73 % to 90 % at a high blood flow rate of 85 mL min^-1^ (Figure 5d) demonstrating a clinically relevant performance during pumpless operation. pO_2_ was significantly enhanced after the LAD and a maximum value of 66 mm Hg was reached at a blood flow rate of 85 mL min^-1^ (Figure 5e). pCO_2_ was also reduced after the LAD (Figure 5f).

The parameters pO_2_, SaO_2_, and pCO_2_ were also measured at the femoral artery and right atrium (mixed deoxygenated blood, venous blood from systemic circulation and oxygenated blood returning from the LAD; ultrasonography of the right atrium of the heart was performed to assure that the catheter was correctly placed as shown in the Supplementary Figure S3) to determine the impact of the LAD on systemic oxygenation for the piglet. Measurements were performed when the piglet was under the hypoxic condition. Both systemic pO_2_ and SaO_2_ (femoral) increased when the LAD exposed to pure oxygen environment was connected with the hypoxic piglet with a maximum increase of 41 mm Hg to 66 mm Hg in pO_2_ and 52 % to 86 % in SaO_2_ (Figure 6a and b). When the LAD was exposed to room air, the increase in systemic pO_2_ and SaO_2_ was lower but an increase of pO_2_ from 42 mm Hg to 63 mm Hg and Sa O_2_ from 56 % to 84 % at a blood flow rate 88 mL min^-1^ was achieved (Figure 6d and e). pCO_2_ was decreased in both oxygen environment and room air (Figure 6c and f).

**Figure 6:**
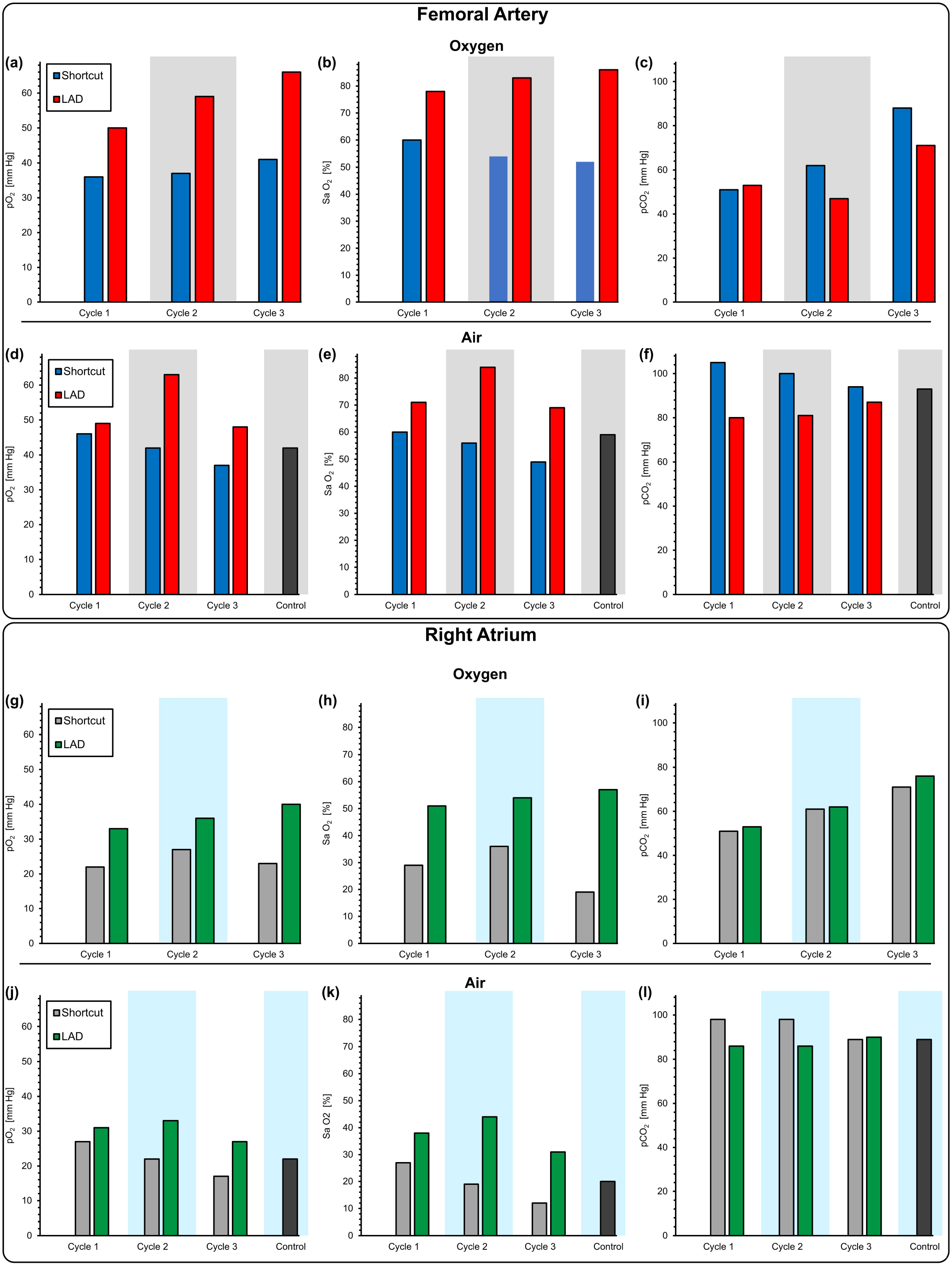
Gas exchange at the femoral artery and the right atrium when connected in-vivo to a piglet. Measurements were of the blood at the femoral artery and the right atrium when the LAD is connected to the piglet and pumped by the arterio-venous pressure difference (a) pO_2_,(b) Sa O_2_, (c) pCO_2_ for femoral artery using oxygen as the sweep gas, (d) pO_2_, (e) Sa O_2_, (f) pCO_2_ for femoral artery exposed directly to room air, and (g) pO_2_, (h) Sa O_2_, (i) pCO_2_ for right atrium using oxygen as the sweep gas, (j) pO_2_, (k) Sa O_2_, (l) pCO_2_ for right atrium exposed directly to room air.

Similar results was observed when measurements were performed at the right heart (atrium). Systemic pO_2_ and Sa O_2_ significantly increased when connected to the LAD but their values at the atrium (Figure 6g and h) were lower than levels immediately after the LAD as blood in the atrium is a mixture of the venous blood from the systemic circulation which is usually rich in carbon dioxide and poor in oxygen with the oxygenated blood from the LAD. During each cycle, pCO_2_ was also elevated after connecting the LAD which was placed under an oxygen-rich hood (Figure 6i). When the LAD was exposed to room air, pO_2_ and Sa O_2_ measured from right atrium also showed an improvement in oxygen content (Figure 6j and k).

pCO_2_ was slightly lower after switching from the shortcut to the LAD extracorporeal connection under hypoxic conditions suggesting that the LAD could be more effective in removing carbon dioxide while exposed to room air (Figure 6l). As expected, the oxygen saturation level dropped in both the femoral artery and the right atrium when the piglet was in hypoxic state and the LAD was inactive (Figure 6). However, the saturation level significantly increased when the LAD was connected. For instance, when exposed to the hypoxic condition the oxygen saturation level in the right atrium dropped to 19 % but increased to 57 % after connecting the LAD (Figure 6h). This effect was observed when the LAD operated in room air with a maximum rise from 31 % to 59 % in Sa O_2_ (Figure 6k).

## Discussion

We tested a new LAD in a newborn piglet model which was constructed of 16 stacked MBOs and designed to support preterm and term infants with respiratory failure. In this study, the LAD provided significant gas exchange throughout a period of ~ 200 minutes. The current extracorporeal circuit achieved blood flow rates through the LAD was on average one-third of the cardiac output which was adequate to maintain the piglet cardiovascularly stable throughout the experiment^[16]^. Our concept of the LAD includes the features of an artificial placenta which allows the newborn to continue to breathe while a partial fetal circulation via the umbilical artery, LAD circuit, and the umbilical vein is established^[16]^. An important element is that the newborn will continue to breathe while the LAD provides additional gas exchange and allows the lungs to heal. In this way, lifelong side effects due to performing mechanical ventilation at the physiological limit could be prevented. Although our concept of the artificial placenta is designed to be connected to the umbilical vessels^[16]^, at this developmental stage, we used the vascular access via carotid artery and jugular vein. This is a compromise of the piglet model. We found that in a one-day-old piglet, the lumen of the umbilical vessels became fibrotic. Using a Seldinger wire technique, only catheters not bigger than an outer diameter of 1.2 mm could be placed which do not provide sufficient blood flow. Other researchers have used lamb models which have umbilical vessels with larger diameters^[1–3,37]^. Also, they reconnected the artificial placenta immediately after birth which prevents umbilical vessels from obliteration. Currently, there are two different approaches for artificial placenta type oxygenation. One concept aims to develop an artificial womb for prolongation of the pregnancy. This would allow the developing infant to maturity while submerged in artificial amniotic fluid. However, the entire gas exchange like in a womb would be facilitated by the artificial lung device^[1,2,37–39]^. Current studies in this version have used commercial, high priming volume, hollow fiber oxygenator and an external pump for blood perfusion. Also, a preterm lamb that matches the appropriate gestational age but are heavier (weight ~ 5 kg) than human neonates (1 to 2 kg) were used. One of the reasons for the use of lambs was the unavailability of small priming volume, pumpless oxygenators of the kind demonstrated in our study.

The second approach which is used in this study, is to provide partial oxygenation support to a newborn neonate in respiratory distress to reduce the load on the lungs and allow it to recover while it continues to breath^[16]^. We specifically designed and optimized our microfluidic LAD for human preterm and term neonate resulting in the choice of a piglet model as it has similar weight, blood volume and physiological characteristics to a human preterm neonate among various animal models. The stackable modular design provides a flexible application from 500 g to 4 kg body weight and facilitate pumpless extracorporeal support with the minimum possible priming volume. Furthermore, the ability to custom fabricate the microfluidic oxygenator allowed us to achieve the specific design criteria for the priming volume, hydraulic resistance and oxygen uptake suitable for human preterm neonates. By using the piglet of the same birthweight as preterm neonates, we were able to accurately test the extracorporeal system for oxygenation performance. The smaller priming volume of this oxygenator also mitigated the effect of hemodilution when the saline that was used to fill it was mixed with the venous blood. The custom and modular design was also critical for pumpless operation as hydraulic resistance of the oxygenator could be tailored to produce adequate blood perfusion and oxygen uptake based on the baby’s weight and an external pump could be eliminated.

Since the arteriovenous pressure difference in a neonate is typically between 20 – 60 mm Hg^[12,16,29–31]^, such a pressure head applied to this LAD would generate a flow rate between 65 – 86 mL min^-1^ in this optimized LAD which meets the requirement of a blood flow rate of 20 – 30 mL min^-1^ kg^-1^ of a neonate typically needed to provide lung assist function (Figure 4. a and e). In this range of flow rates and under in-vitro conditions, this LAD was able to increase the oxygen saturation level from ~ 75 % to ~ 100 % which would be sufficient to fulfill oxygenation need for these preterm babies^[12,16,29–31]^.

Of note, the hemoglobin of the piglet was at a lower level (hematocrit: 23 % to 27 %) in these experiments. A preterm infant typically has considerably higher hematocrit level (40 to 60%)^[40]^, and as a result, the oxygen-carrying capacity of the blood would allow higher amounts of gas exchange and oxygen delivery compared with the demonstrated capacity in this study. The performance of this LAD connected to human neonate can be estimated for comparison purposes by calculating its performance at these higher hematocrit levels. Such calculation (as shown in the Supplementary section, Figure S7 and Figure S8) should include physiological trends including post-natal decrease in hemoglobin content^[40]^ and corresponding increase in systemic blood flow rate ^[41]^ during first days of life for which measurements exist. The estimation indicates that for preterm neonates with weights from 500 g to 1 kg, over a period of 96 hours after birth, the current configuration of the LAD is capable of providing a gas exchange capacity of 4.2 mL min^-1^ kg^-1^ using oxygen as the sweep gas or 3.1 mL min^-1^ kg^-1^ while the LAD would be exposed to room air. This corresponds to ~ 50 – 75 % (~ 7 mL min^-1^ kg^-1^) of the total oxygen requirement for a resting 1 kg preterm infant at 96 hours postnatal age^[42,43]^. This capacity far exceeds that required for our artificial placenta type oxygenator which is designed to assist the lungs and provide ~ 30 % of this total oxygen requirement^[8,16]^.

The arteriovenous connection used in our approach, directs partially oxygenated blood from the lungs into the LAD. The LAD assists by further increasing the oxygen saturation to 100% before returning it back to the body. Calculations show that the LAD is capable of supporting a 1.5 kg neonate by increasing the oxygen saturation of 50 mL min^-1^ of its blood from ~75% to 100% in room air thereby providing the 30% of the total oxygen requirement and preventing respiratory distress. Furthermore, the return of oxygenated blood to the right heart also assists in counteracting hypoxic pulmonary vasoconstriction and decrease pulmonary resistance and improve lung perfusion^[44]^.

The piglet was cardiovascularly stable throughout the entire in-vivo experimentation that occurred over a period of ~ 200 minutes. We did not observe bradycardia or tachycardia, or arrhythmia and the blood pressure was in the desired range. Our LAD was connected in parallel to the systemic circulation and considerable amounts of blood bypassing the systemic circulation extracorporeally. The bypassed blood flow (65 – 86 mL min^-1^) reaches up to one-third of the cardiac output. This clinical condition is similar to a patent ductus arteriosus or a ventricle septum defect and could pose a risk to the heart of high cardiac output failure. Clinical signs would be tachycardia, low systemic blood pressure, and lactacidemia. We found that blood pressure of the piglet remained stable (Figure 4. d and h), no significant increase of the heart rate (**Error! Reference source not found.**. b and f) were noted, and no metabolic acidosis could be observed (Figure S6 in the supplementary).

The LAD was able to increase the oxygen saturation of blood to desired target levels for preterm neonates with respiratory failure. Systemic oxygen levels were measured using a pulse oximeter. A typical range of SpO_2_ is 85 to 95% for normal neonates, preferably above 90%^[45]^.

To achieve hypoxic condition, we reduced the ventilation rate of the paralyzed piglet and targeted hypoxic SpO_2_ of 60%. It was expected that adding the LAD will increase the systemic SpO_2_ to a range of 85 – 95%. Notably, this target range for SpO_2_ was achieved in all three cycles when the ventilation gas for the LAD was pure oxygen (results are shown in Figure S5 in the supplementary). Of particular note was in the third cycle, where the SpO_2_ was increased from 57 % (under hypoxic condition) to 91% (upon connection with the LAD). When room air was used as the ventilation gas, the LAD was still able to increase SpO_2_ during all cycles. However, the SpO_2_ does not reach above 85% except in the second cycle where it increased from 69 % to 90 % upon the introduction of the LAD). This is because of the excessively low oxygen saturation (42% in cycle 1) induced in the hypoxic condition in cycles 1,3. Even then, the LAD was able to increase systemic levels of SpO_2_ to ~ 80% which approaches the systemic SpO_2_ in healthy newborn piglets with a weight of 1.77 ± 0.13 kg varies from 84 to 96 %^[46]^. Overall, the in-vivo experiments demonstrate that the LAD could significantly improve systemic levels of SpO_2_ either using oxygen or room air as the ventilation gas in a piglet under hypoxic condition.

The piglet required continuous normal saline infusion at a high rate of more than 15 ml/kg/hr due to significant insensible water losses. There are two main contributing factors: first, the piglet is exposed to a radiant warmer to maintain a body temperature of 39°C which promotes evaporation of water. Studies of ventilated ill newborns under a radiant warmer showed that these infants lose 28.04 · *e*^−1.73(*Weight in kg*)^ ml/kg/hr^[47]^. Second, the LAD is composed of PDMS which is permeable to water. Water from the circulating blood could have evaporated as a result. However, fluid support during the experiments prevented signs of metabolic acidosis, the hydrogen carbonate remained in desired range, as well as the urine output was stable indicating sufficient hydration. As a result, a possible future modification to minimize evaporation and prevent heat loss can be to surround the LAD by humidified air.

We used low dose hydrocortisone and dopamine in this study to allow sufficient stress response, to minimize inflammatory responses and to provide vascular tone. In our experimental setting, the piglet underwent surgery for the establishment of the tracheostomy and catheters were placed invasively. Further, exposing the piglet’s blood to the extracorporeal tubes and LAD can potentially induce inflammation including mild capillary leak syndrome. An increase in vascular tone by dopamine as well as the effects of hydrocortisone on the reduction of inflammation and blood pressure provide stable conditions for this experiment.

An essential characteristic of the LAD is hemocompatibility because the LAD should not induce hemolysis or blood clot formation. The hematocrit of the piglet remained throughout the experiment at expected levels. There was only a marginal drop of the hematocrit by 4 % from 27 to 23 %. We estimated the blood loss to be approximately 4 mL/kg, which could be explained by several blood collection during the study. No side effects of a clinically significant anemia such as tachycardia or acidosis could be recognized. However, future studies should measure hemolysis.

To prevent blood clotting and to establish hemocompatibility, a key feature of our LAD is the coating of its inner surfaces. In vitro, we performed several experiments to show that heparin was adsorbed on PDMS surfaces and was fairly stable (It could be hypothesized that some of the heparin was absorbed by the PDMS surfaces). This bonding could be explained by the formation of hydrogen bonding or electrostatic interaction between heparin molecules and PDMS chains^[34]^. We also stored the plasma samples after performing the plasma clotting assays observing if plasma would be clotted after being exposed to an external surface without heparin coating. We observed that the plasma samples were not clotted even after being exposed to a non-coated surface for few hours suggesting that the heparin was not covalently bonded to PDMS. This was expected and was also reported in another study^[34]^. Therefore, in our experiment, we primed the LAD with a normal saline and heparin solution. However, this approach alone was not sufficient to prevent blood clotting and we used it in conjunction with a typical systemic dose of heparin for systemic anticoagulation. Nevertheless, coating was important in preventing any deleterious clot formation on the surfaces of the oxygenators and critical for smooth operation of the LAD in in-vivo experiments. In preterm infants, this approach of systemic anticoagulation may not be feasible. Preterm infants are at high risk for intraventricular hemorrhage (IVH); in particular, those infants who are born preterm from mothers with chorioamnionitis, HELLP, or preeclampsia. Systemic anticoagulation would elevate this risk significantly. Initial work to achieve hemocompatibility of the LAD has been done by our group. The coating of inner surfaces of the LAD, the tubes, and the catheter with the novel antithrombin-heparin-complex are promising^[36,48]^. This type of coating could be applied to all internal surfaces to enhance hemocompatibility.

## Conclusion

In this study, a pumpless LAD using microfluidic blood oxygenators for preterm neonates suffering from respiratory failure was developed. In vitro assessments were done by blood to investigate the gas exchange capacity of this LAD resulting in an acceptable gas transfer in room air and ample oxygenation in an oxygen-rich environment. A simple coating technique was developed to coat blood-in-contact surfaces. The functionality and stability of this coating were evaluated in vitro suggesting that the coating was effective and stable enough to be used in further animal studies. In the next step, the LAD was tested in a pumpless operation connected to a newborn piglet. Results from the piglet experiments revealed the effectiveness of this LAD in gas exchange without complications and agreed with in vitro results. Further steps in the development of the LAD include the installation of hemocompatibility, biocompatibility, and developing via the umbilical vessels.

## Methods

### The design of microfluidic blood oxygenator device

The microfluidic blood oxygenator (MBO) was constructed entirely out of polydimethylsiloxane (PDMS) comprising three components: (i) a blood vascular network that spreads the incoming blood into a thin uniform layer suitable for fast gas exchange, enclosed by (ii) two thin composite membranes made of polytetrafluoroethylene (PTFE) and PDMS with high permeability for blood gases and sufficient stiffness to withstand the physiological pressure encountered, and (iii) a tapered shaped inlet/outlet to access the blood vascular network. The blood vascular network was a microchamber with a height of 180 μm (or 130 μm) containing an array of 1 x 1 mm square-shaped micropillars. The design which is shown in Figure 7a was optimized to ensure that a uniform blood distribution was achieved. It had a surface area of 29.19 cm^2^, and a priming volume of 0.52 mL. Tapered inlet and outlet were integrated on top of the blood vascular network to ensure smooth distribution of blood from a cylindrical inlet connection into the vascular network (Figure 7b). The design minimizes shear stress that could lead to hemolysis, platelet activation, and clot formation. A numerical modelling was conducted by COMSOL Multiphysics (Version, Producer, country) to show the uniform flow distribution in the blood vascular network without creating any dead zone in the inlet (Figure 7c).

**Figure 7:**
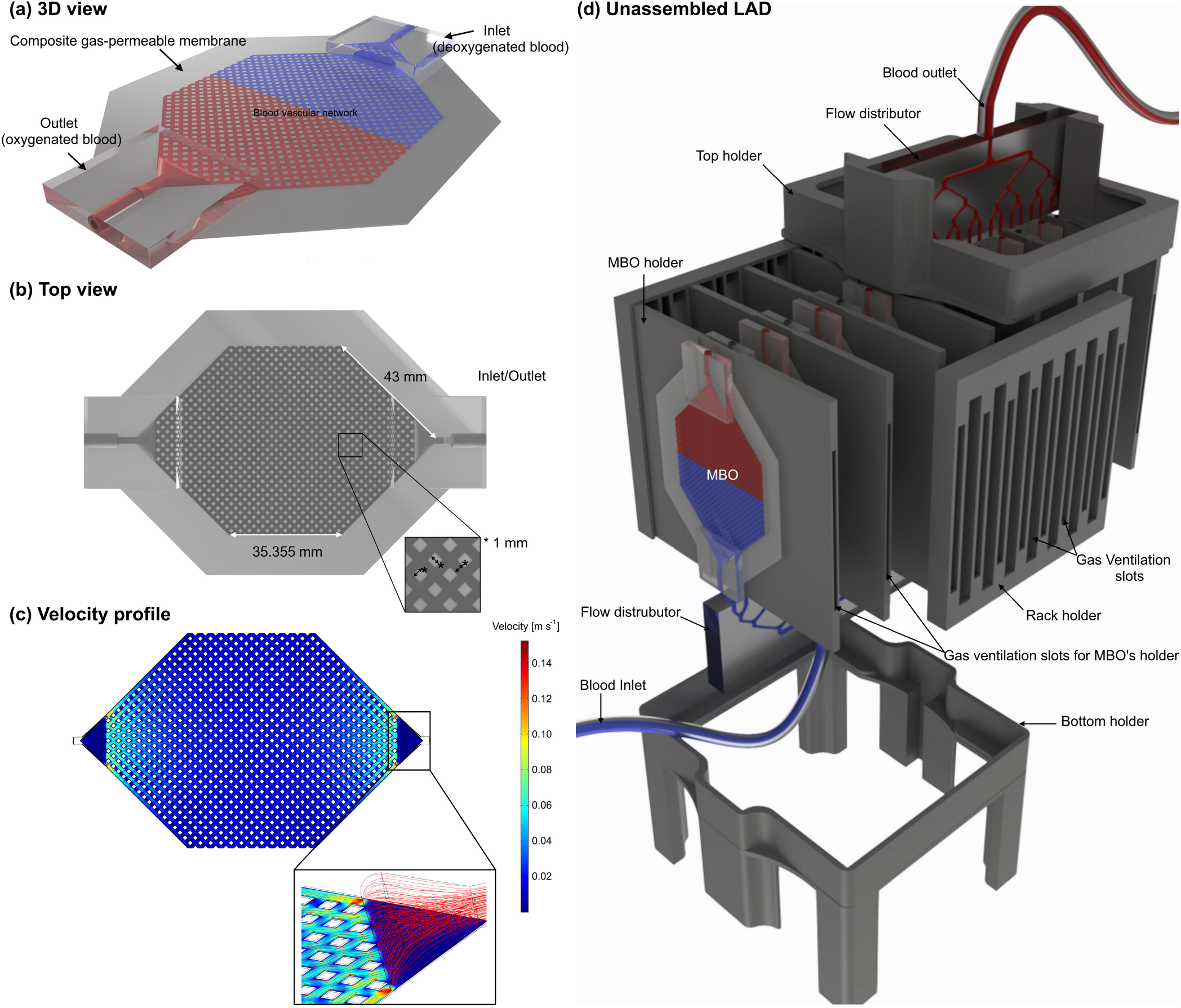
(a) 3D schematic drawing of MBO with tapered inlet/outlet configuration, (b) top view of MBO, (c) the velocity profile of MBO at a blood flow rate of 5 mL min^-1^ showing a uniform distribution with a gentle change in flow direction and no dead zone, and (d) 3D drawing of the LAD in unassembled format showing all 3D-printed components, MBOs, and flow distributors. There are fixed gaps among all MBOs for air to flow convectively. Blood is fed to the LAD from the bottom flow distributor and leaves the LAD from the top one to ensure that no air bubble is trapped inside.

### MBO fabrication

A precisely controlled layer-by-layer microfabrication process was developed to manufacture all MBOs (Figure 8a – e) and described in previous publication^[30]^. Briefly, a negative mold of MBOs (Figure 8a) was fabricated by conventional photolithography using a negative photoresist (SU8 3035, MicroChem. Corp., Westborough, MA, USA). PDMS (Sylgard 184, Dow Corning, Midland, MI) with a ratio of 10:1 (base to curing agent) was prepared, and coated on the mold using a spin-coater with a speed of 750 RPM for 60 s, followed by being cured on a hotplate at 85 °C for 30 minutes. Another layer of PDMS was spin-coated (2000 RPM for 40 s) and a porous PTFE membrane (pore size of 1 μm and a porosity of 83%, PTU103001H, Sterlitech, Kent, WA, US) was embedded into the wet PDMS, and cured in an oven at 85 °C (Figure 8b). The blood vascular network was removed from the mold (Figure 8c). Another porous hydrophobic PTFE membrane (pore size of 0.22 μm and a porosity of up to 90 %, PTU023001, Sterlitech, Kent, WA, US0) was fixed on a hydrophobic substrate, which was made by sticking a piece of PTFE film on a silicon wafer, and covered by wet PDMS for ~ 15 minutes to ensure that all pores were filled by PDMS. This substrate with PDMS-wetted PTFE membrane was spun at 8000 RPM for 60 s and brought into contact with the blood vascular network (Figure 8d). The device was placed on a leveled surface and cured overnight at room temperature. Two inlets with tarpapered configuration were placed on their designated locations on the top of the blood vascular network being glued by half-cured PDMS, and they were cured in an oven at 85 °C for an hour (Figure 8e). A scalpel and a biopsy puncher were used to remove residual PDMS inside the inlet and outlet and the other sides were sealed by a piece of cured PDMS. Figure 8f shows an MBO with the tapered inlet which was filled with dyed DI water.

**Figure 8:**
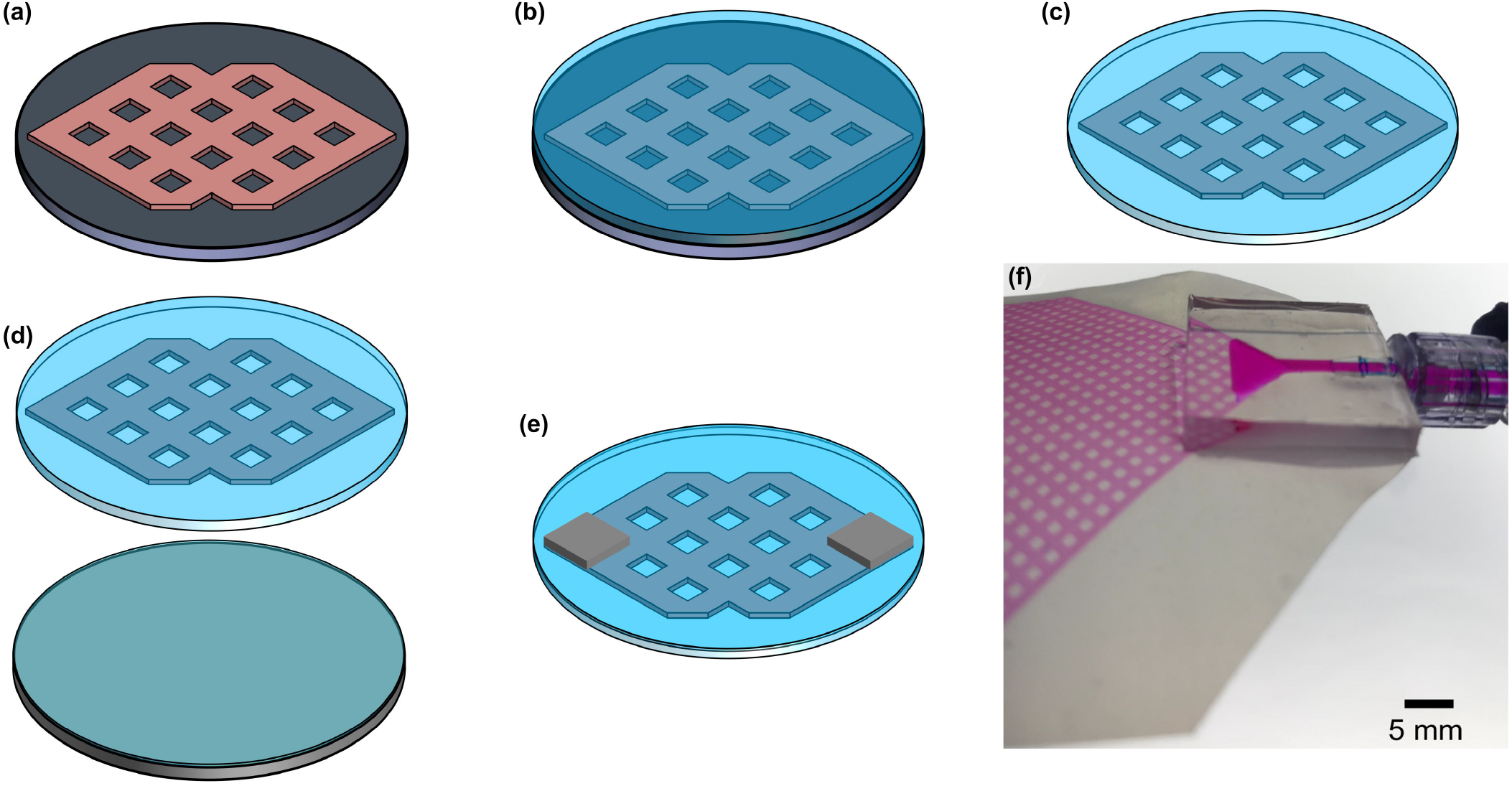
(a – e) the microfabrication process of MBOs and (f) a close-view image of an MBO with the tapered inlet configuration filled with dyed DI water.

### The assembly of the compact neonatal lung assist device

A single MBO cannot provide enough oxygenation to support preterm neonates with respiratory failure with different weights. As a result, the lung assist device (LAD) was constructed by assembling 16 MBOs in a parallel configuration (Figure 7d). The frame was 3D-printed to arrange the MBOs in a compact form and consisted of several fixtures to hold MBOs along with a top and bottom holder to maintain fixed gaps between the MBO holders. Each MBO holder was designed to hold two MBOs at a fixed distance apart from each other to facilitate convective gas flow on all sides of the MBO, naturally. Then, eight such MBO holders were inserted into the appropriate slots in the top and bottom holders to form the compact LAD.

Autodesk Inventor software (2018, USA) was used to draw all components for the compact LAD. All holders were manufactured by a Prusa i3 mk3 3D printer (Partyzánská, Praha Czech Republic) including 8 MBO holders, one top and bottom holder, two rack holders, and two flow distributor holders (the bottom one was 3D-printed by four legs). The flow distributor was casted out of a 3D-printed mold, which was made by a Formlab Form 2 3D printer (Somerville, MA, USA) using PDMS with the similar mixing ratio and bonded later to a cured 1-mm-thick PDMS by means of a flame treatment^[49]^ and an oxygen plasma activation (1.5 minutes exposure, an oxygen pressure of 900 mm Torr, Harrick plasma cleaner, Ithaca, NY, USA). For each LAD, at least 32 MBOs were fabricated and their hydraulic pressure characteristics were quantified. Then, those with similar hydraulic behavior were chosen for LAD assembly.

### In vitro testing of LAD using bovine blood

Bovine blood was purchased from Lampire Biologicals Laboratories (Bovine 7200807-1L, Pipersville, PA) and heparinized by a concentration of 3 units mL^-1^ right away after delivery and was stored in a fridge overnight. A hollow fiber oxygenator (PDMSXA-1.0, PermSelect VR, Ann Arbor, MI, USA) was used to lower the oxygen content of blood to ~ 60 % by passing a carbon dioxide/nitrogen gas mixture (5%/95% v/v) into the shell while blood was fed into the fibers using a peristaltic pump (Ismatec model ISM832C, Ismatec, Glattbrugg, Switzerland) at 15 – 25 ml min^-1^. The blood was collected in a sealed bottle and stored again in a fridge overnight. Prior to each experiment, the bottle of the blood with a sealed cap was warmed up to room temperature and an open-loop circuit was created by pumping (a peristaltic pump was placed in the circuit) the blood from the bottle to the LAD, then to a waste bottle. Blood samples were taken before and after the LAD and analyzed by a point-of-care blood gas analyzer (GEM Premier 3000, Instrumentation Laboratory, Lexington, MA, USA). A Complete MicroHematocrit System (StatSpin CritSpin, Norwood, MA, USA) was used to measure the hematocrit level of blood samples as well.

### Heparin coating

Heparin sodium injection USP with 1000 USP Units per mL concentration was a high molecular weight polysaccharide and was purchased from Sandoz Canada Inc. It was a sterile, pyrogen-free solution with high purity derived from porcine intestinal mucosa.. FITC-labeled heparin was obtained from Creative PEGWorks (HP-201, Chapel Hill, NC, USA) and was also a high molecular weight polysaccharide (27k). FITC-labeled heparin was only used for visualizing the quality of coating inside MBOs. Heparin sodium injection USP was used for coating LADs in the animal experiments as well as in the in-vivo experiment for systemic heparinization. Generally for the coating, heparin sodium injection USP or FITC-labeled heparin was circulated in the LADs or MBOs prior to each experiment in a closed-loop configuration for 4 to 5 hours followed by washing with PBS or normal saline (only for LADs tested in the animal experiments) to remove any uncoated heparin for ~ 15 minutes.

### FITC-fibrinogen adhesion test

Human Fibrinogen Purified FITC Labeled was obtained from Innovative Research Inc (IHUFBGFITC50MG, Novi, MI, USA) and stored at −20 °C. Prior to each experiment, FITC-fibrinogen was warmed up to room temperature and was added to PBS buffer with a pH of 7.4 to investigate the antithrombotic property of the noncoated and coated surfaces. Coated and non-coated MBOs were perfused for 3 hours in separate closed-loops containing 2 mg mL^-1^ FITC-fibrinogen (a total volume of ~ 5 mL). and the MBOs were washed with PBS (pH = 7.4) for 30 minutes with a flow rate of 3 mL min^-1^. All other components used in the loop for testing MBOs such as tubes and connectors were coated with heparin using the same procedure. This was done to ensure that no part of the circuit would induce clot formation.

### Plasma clotting assays in MBOs

Human plasma was purchased from Innovative Research Inc (Novi, MI, USA) and prepared by mixing 5 mL of 25 mM CaCl_2_ with 5 mL human plasma^[50]^. The recalcified plasma was transferred immediately to a closed test loop containing the MBO. Care was taken to avoid the formation of air bubbles. The plasma was then circulated in the loop and clotting was assessed by measuring the pressure change at the inlet of the oxygenator.

### Experimental setup of the newborn piglet model

The newborn piglet was first weighed (1.78 kg) and anesthetized by intraperitoneal injection of sodium pentobarbital (30 mg kg^1^, MTC Pharmaceutics, Cambridge, ON, Canada). To monitor the body temperature and maintain it at 39 °C during the experiment, an infant servo control (ISC) probe was placed on the piglet’s abdomen. To provide controlled mechanical ventilation, an endotracheal tube (3.5 mm inner diameter, Portex, Keene, NH, USA) was placed via tracheostomy and connected to a Servo 300 ventilator (Siemens, Mississauga, ON, Canada). The ventilator was adjusted in a way to assure that the newborn piglet experienced a physiologic gas exchange at the beginning: peak inspiratory pressure (PIP) of 14 cm H_2_O, positive end-expiratory pressure (PEEP) of 5 cm H_2_O, inspiratory time (IT) of 0.33 s, and respiratory rate (RR) of 31 breaths min^-1^. A mixture of oxygen (30 %) and nitrogen (70 %) was humidified and warmed up to 38 °C for delivering to the piglet (at a flow rate of 8 L min ^1^). 22-G angio-catheters were used to cannulate femoral and abdominal veins for maintenance fluids: sodium pentobarbital (16 mg/kg), 5% dextrose (80 mg kg^-1^ day^-1^), hydrocortisone and dopamine infusion (2 μg kg^-1^ min^-1^). The dopamine was adjusted (2 to 20 μg kg^-1^ min^-1^) to maintain the mean systematic blood pressure in a range of 68 – 80 mmHg. During the experiment, the piglet was paralyzed using a muscle relaxant (Pancuronium) with an initial bolus 0.2 mg kg^-1^ and continuous infusion of 20 μg kg^-1^ min^-1^.

Two 14-G 1.1-inch angio-catheters (Angiocath) were used to cannulate left carotid artery and right jugular vein for creating an extracorporeal circuit in which the LAD was attached. This circuit consisted of two loops: 1) shortcut bypass and 2) the LAD bypass. Once one of the bypasses was opened, the other one was closed. A flow sensor (in the main line) with a monitor and a pressure transducer (in a branching line connected to the main line) with a blood pressure monitor were installed in the circuit to report the extracorporeal blood flow rate and blood pressure.

While the extracorporeal circuit was connected, a bolus of 400 units kg^-1^ of heparin as an anticoagulant was delivered to the piglet and maintained at an infusion rate of 30 units kg^-1^ h^-1^. Access to the right femoral artery was achieved by insertion of a 3.5 Fr Argyle umbilical catheter. This vascular access was used to measure systemic blood pressure, heart rate, and for blood collections. The catheter was perfused by a heparinized normal saline solution with a concentration of 3 units mL^-1^ at a perfusion rate of 1.5 mL h^-1^. A pressure transducer with a blood pressure monitor was attached to the catheter to monitor systemic blood pressure during the experiment. Another vascular access was established via right internal jugular vein using a 3.5 Fr Argyle umbilical catheter which was advanced under ultrasound into the right atrium of the heart which allowed measuring the mixed blood (systemic venous blood and LAD oxygenized). Peripheral oxygen saturation was also monitored using a pulse oximeter which was placed on one of the feet.

The study protocol was approved by the McMaster University Animal Research Ethics Board (AREB #10-03-14). The animal was humanely euthanized with Euthanyl Forte (Bimeda-MTC Animal Health Inc., Cambridge, Ontario, Canada) as approved in the animal utilization protocol.

## Acknowledgments

This work was supported by the Natural Sciences and Engineering Research Council of Canada (NSERC) and Canadian Institutes for Health Research (CIHR) through the Collaborative Health Research Program. PRS also acknowledges support from the Canada Research Chairs Program as well as the Discovery Accelerator Supplement grant. CF acknowledges support from Jack Sinclair Research Chair position at McMaster University. The authors would like to thank Devon Jones, a graduate student in CAMEF lab, for her assistance in the animal experiments.

## Additional information

The supplementary materials are available.

