## Supplementary data for "A Pumpless Microfluidic Neonatal Lung Assist Device for Support of Preterm Neonates in Respiratory Distress"

### In vitro carbon dioxide release for LAD

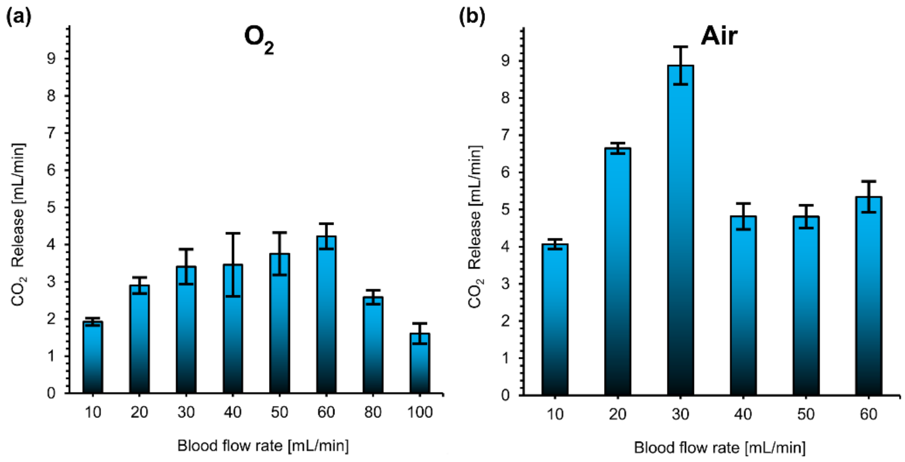


*Figure S1: (a) $CO_2$ release at various blood flow rates while the LAD was consuming oxygen as the sweep gas and (b) $CO_2$ release at various blood flow rates while the LAD was exposed to room air.*

### Sequence of conditions for in vivo study

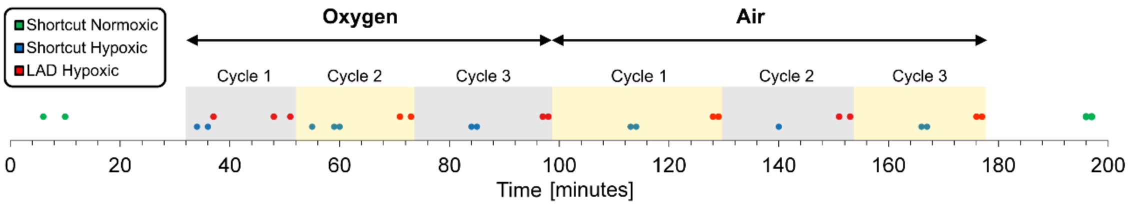


*Figure S2: Time sequence of in-vivo measurements indicating the conditions that the piglet was exposed to and the time points when blood samples were extracted. The LAD was tested first with pure oxygen and then switched to room air. The x-axis represents the time and points represents when blood samples were taken with alternating backgrounds color for cycles changes.*

#### Access to the right atrium of the heart

A 3.5 Fr Argyle umbilical catheter was used to access the right atrium of the heart via the right internal jugular vein. To guide the catheter to the designated location and ensure that the catheter is not clogged, ultrasonography was used.

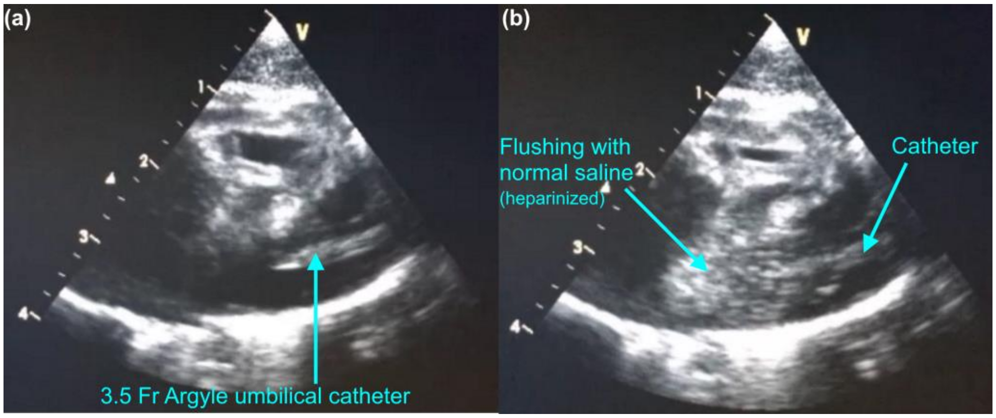


*Figure S3: Ultrasonography of the right atrium of the heart: (a) securing a 3.5 Fr Argyle umbilical catheter and (b) flushing the line with heparinized normal saline solution.*

Figure S3a shows that the catheter was placed in the right atrium of the heart. And it was flushed with heparinized normal saline showing that it was not clogged (Figure S3b).

#### Achieved blood flow rates and pressure drops over time

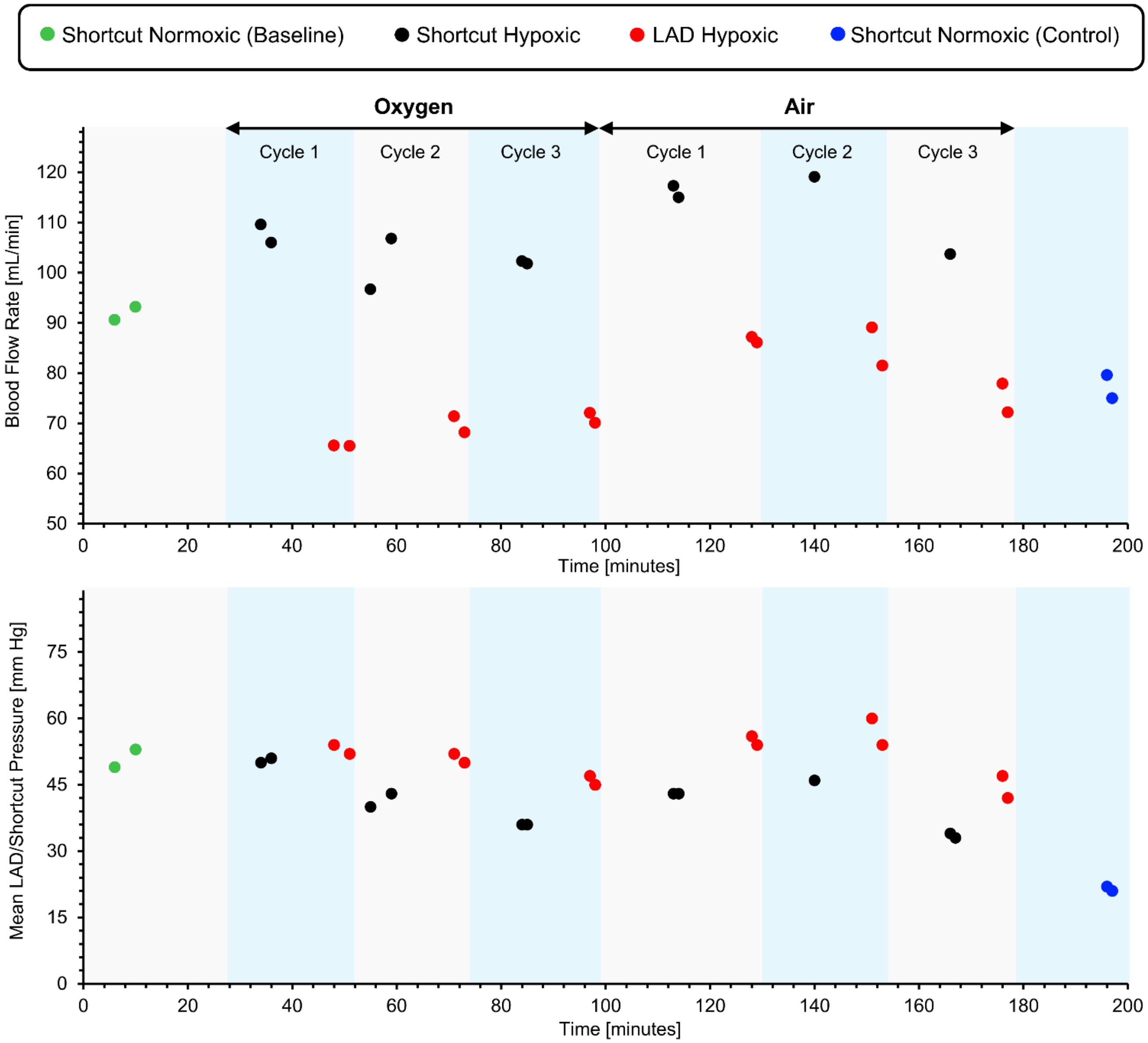


*Figure S4: (a) measured blood flow rates over the period of the experiment and (b) the shortcut or the LAD's pressure during the experiment.*

#### Systemic oxygen saturation measured by pulse oximetry

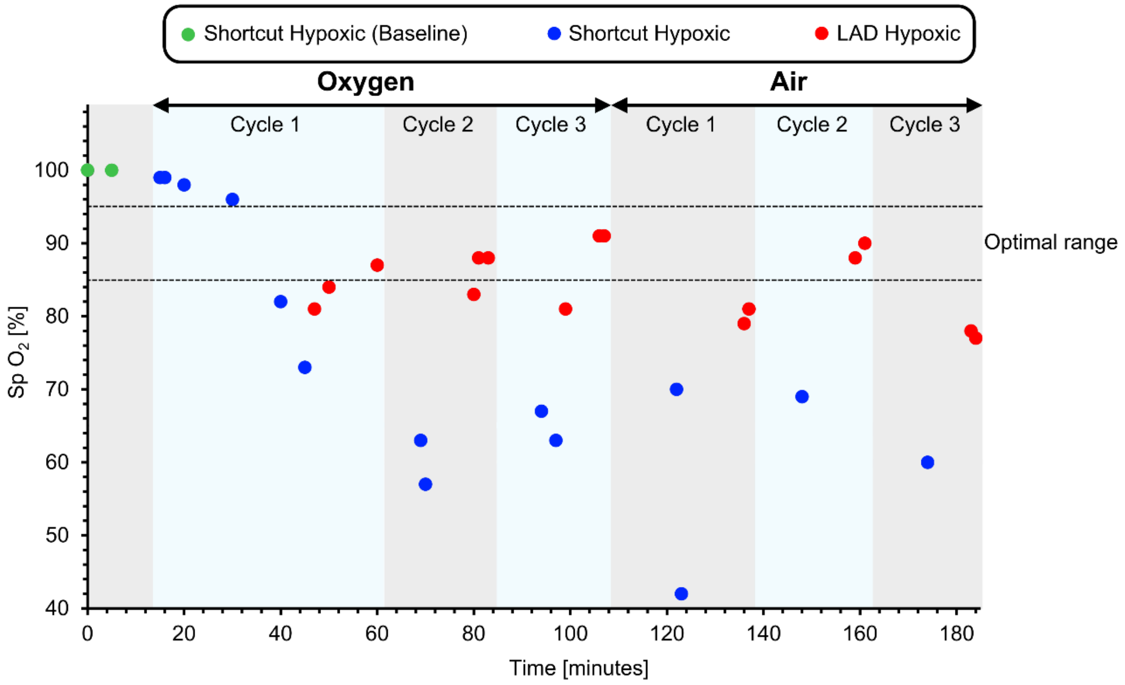


*Figure S5: systemic oxygen saturation level measured by a pulse oximeter placed on the left foot.*

### Metabolic analysis by sampling from the femoral artery

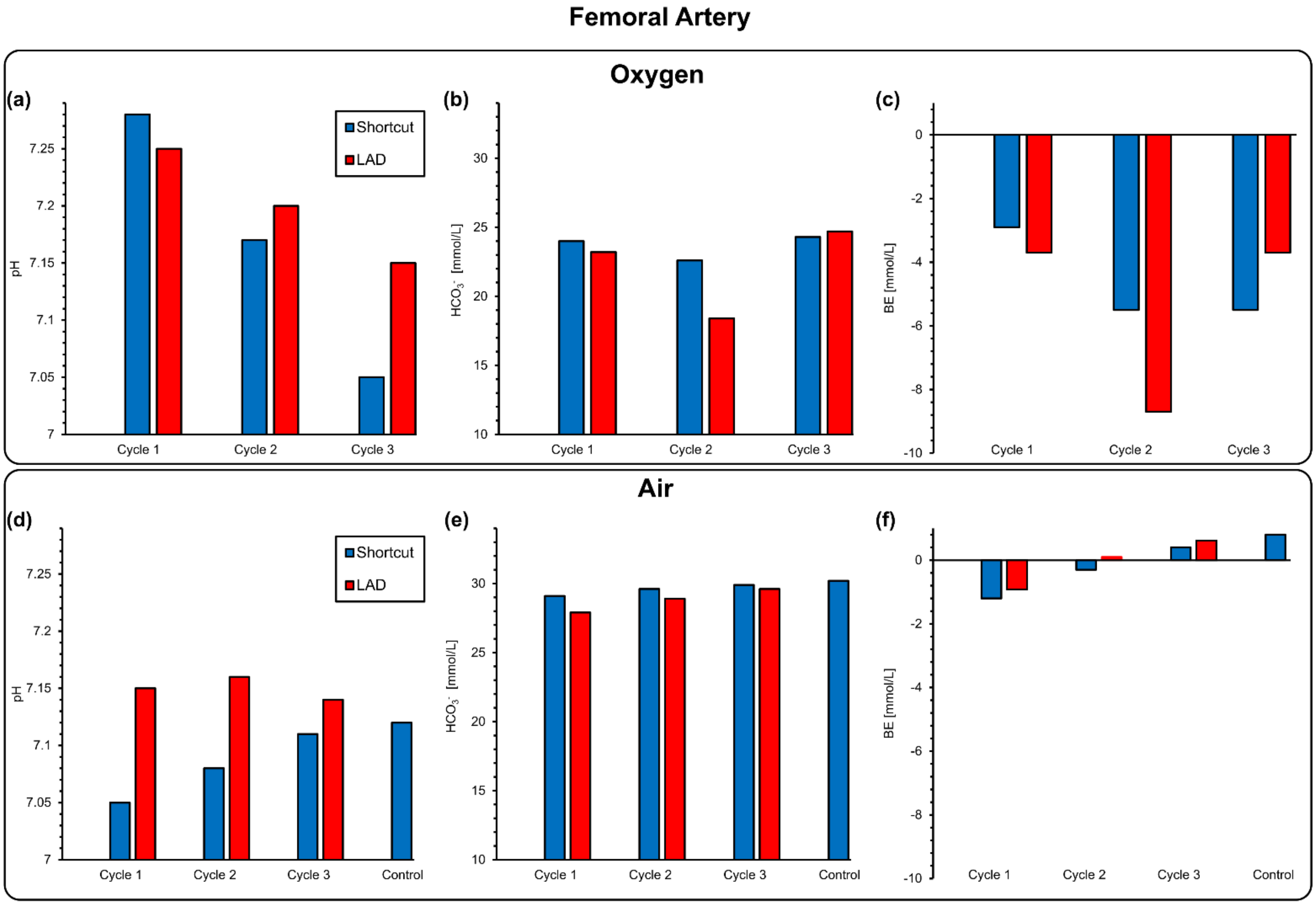


*Figure S6: The metabolic analysis: (a) pH measurement, (b) bicarbonate content, (c) base excess $pCO_2$ for femoral artery using oxygen as the sweep gas, and (d) pH measurement, (e) bicarbonate content, and (f) base excess for femoral artery exposed directly to room air.*

#### Estimated oxygen transfer by the LAD

In this study, the piglet had a lower hematocrit levels compared to preterm neonates. Therefore, to evaluate the real capacity of the developed LAD, we estimated the amount of oxygen transfer for different weights up 1000 grams for both oxygen and air as the sweep gas.

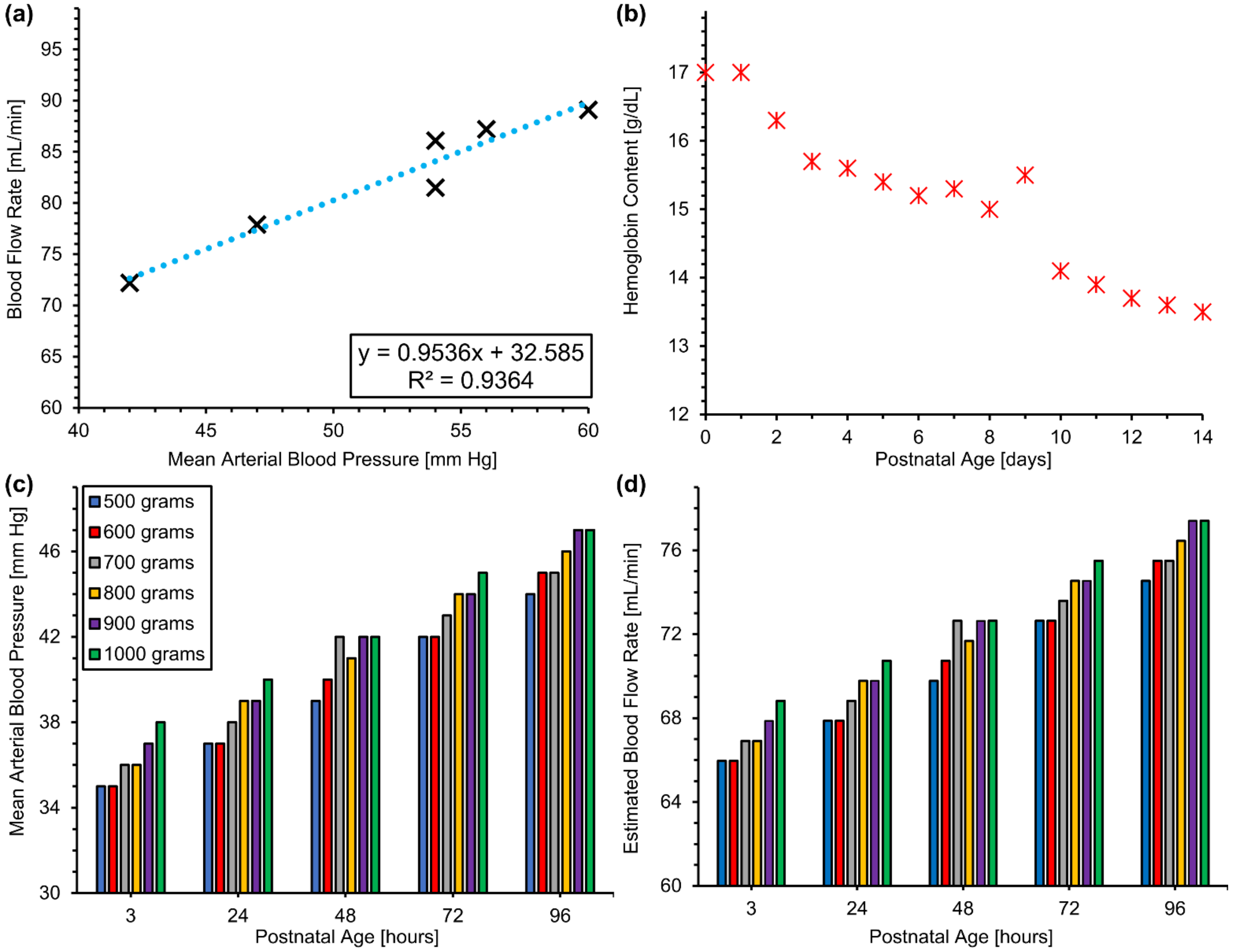


*Figure S7:(a) measured blood flow rates versus mean arterial blood pressure in the piglet experiment, (b) measured hemoglobin contents for preterm infants with a gestational age of 29 – 34 weeks*[1]*, (c) measured mean arterial blood pressure for newborns with different weights over 96 hours after birth*[2]*, and (c) estimated blood flow rates based.*

First, the correlation between mean arterial blood pressure (MABP) and blood flow rates (BFR) were found using the data gathered from the piglet experiment (Figure Sa). A linear relationship between MABP and BFR was established which can be used to estimate the BFR for other MABP values. As hematocrit level in human infants are higher, we used the measured values by Jopling et al.[1]. They measured the hematocrit level for neonates at different gestational age (GA) from 22 weeks to 42 weeks. Data used in this calculation belonged to the group with a GA of 29 – 34 weeks (Figure Sb). As seen here, hematocrit level decreases after birth which would affect the overall blood oxygenation. Moreover, MABP values for different postnatal age (up to 96 hours after birth) and various weights (500 – 1000 grams) were used from the study conducted by Jones et. al.[2]

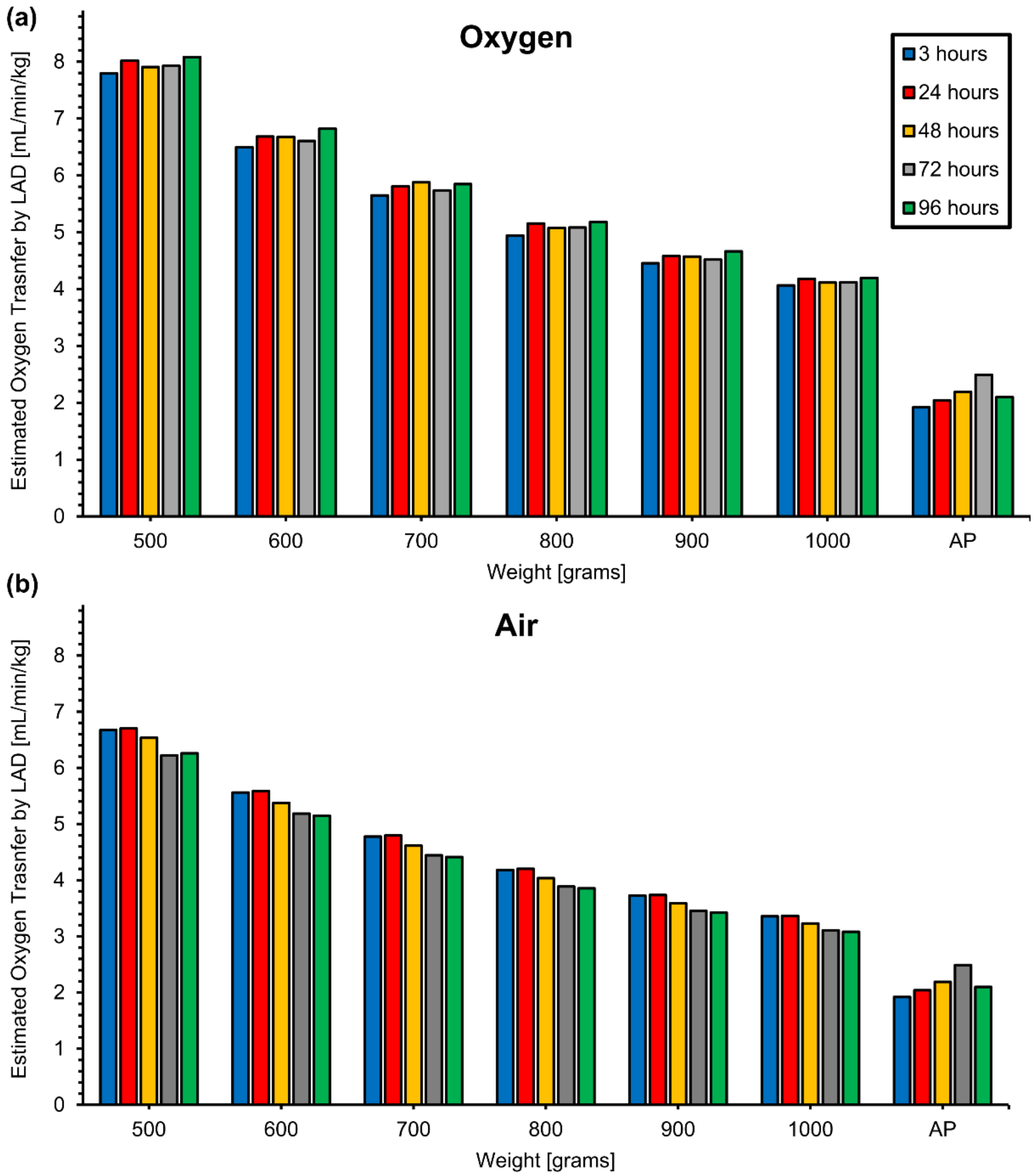


*Figure S8: (a) estimated oxygen transfer by the LAD while consuming oxygen as the sweep gas for preterm neonates with different birth-weights over 96 hours after births and (b) estimated oxygen transfer by the LAD while the LAD is exposed to room air for preterm neonates with different birth-weights over 96 hours after births. AP shows the reference values for different weights in an artificial placenta configuration when the LAD is supposed to provide 30 % of oxygen consumption*[3].

Then, the blood flow rates for each weight was estimated using these MBAP values. It was assumed that the initial value of oxygen saturation level was 75 % for an artificial placenta application and it would be raised to 100 % in an ideal scenario. This means that the maximum increase in oxygen saturation level would be 25 %. Next, the amount of increase in oxygen saturation level was estimated for each flow rate using in vitro data. For instance, the oxygen saturation level after the LAD would be always 100 % for all flow rates while oxygen is assumed to be used as the sweep gas. Under room air condition, the increase in oxygen saturation level was 25.7 % and 23 % at blood flow rates of 50 and 60 mL min$^{-1}$. Finally, the amount of oxygen transfer was calculated for different hours and weights considering the fact the hematocrit level decreases and blood flow rate increases while the baby is growing.

Also, it was assumed that the LAD would provide 30 % of the oxygen consumption which corresponding values are shown in Figure S as references (AP). Based on this calculation, the LAD would be able to provide the required oxygenation in an artificial placenta configuration while using oxygen or air as the sweep gas.

### In vitro comparison between LADs with different channel heights

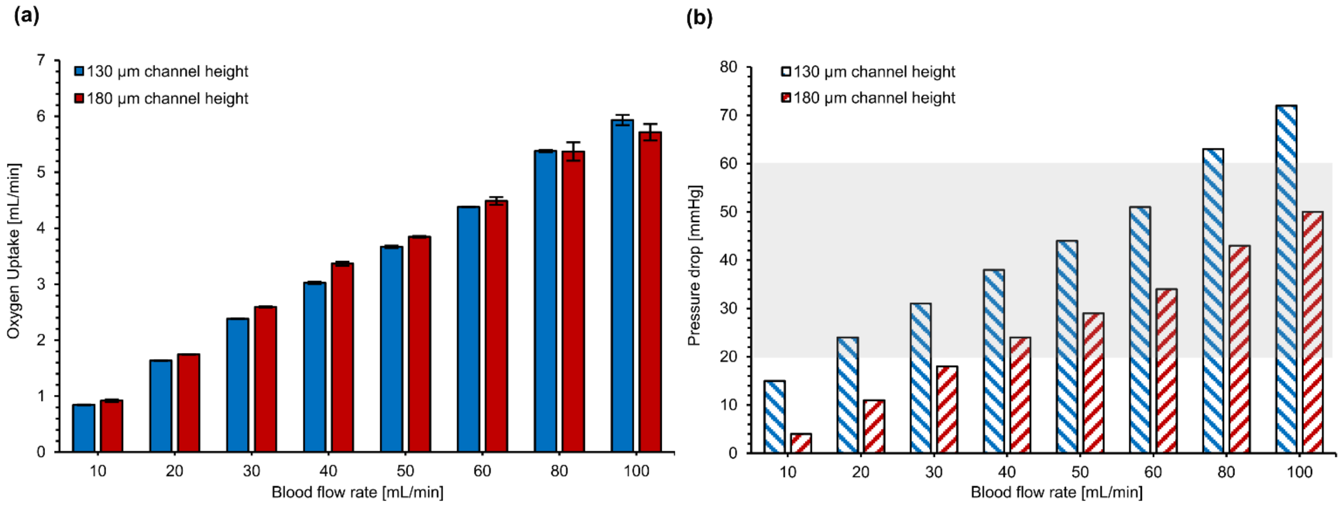


*Figure S9: In vitro comparison between LADs with two different microchannel heights of 130 µm and 180 µm with respect to blood flow rate: (a) oxygen uptake in an oxygen-rich environment and (b) pressure drop (the shaded gray region represents the operating pressure drop range for preterm infants).*

### Cardiovascular parameters for the LAD with a microchannel height of 130 μm

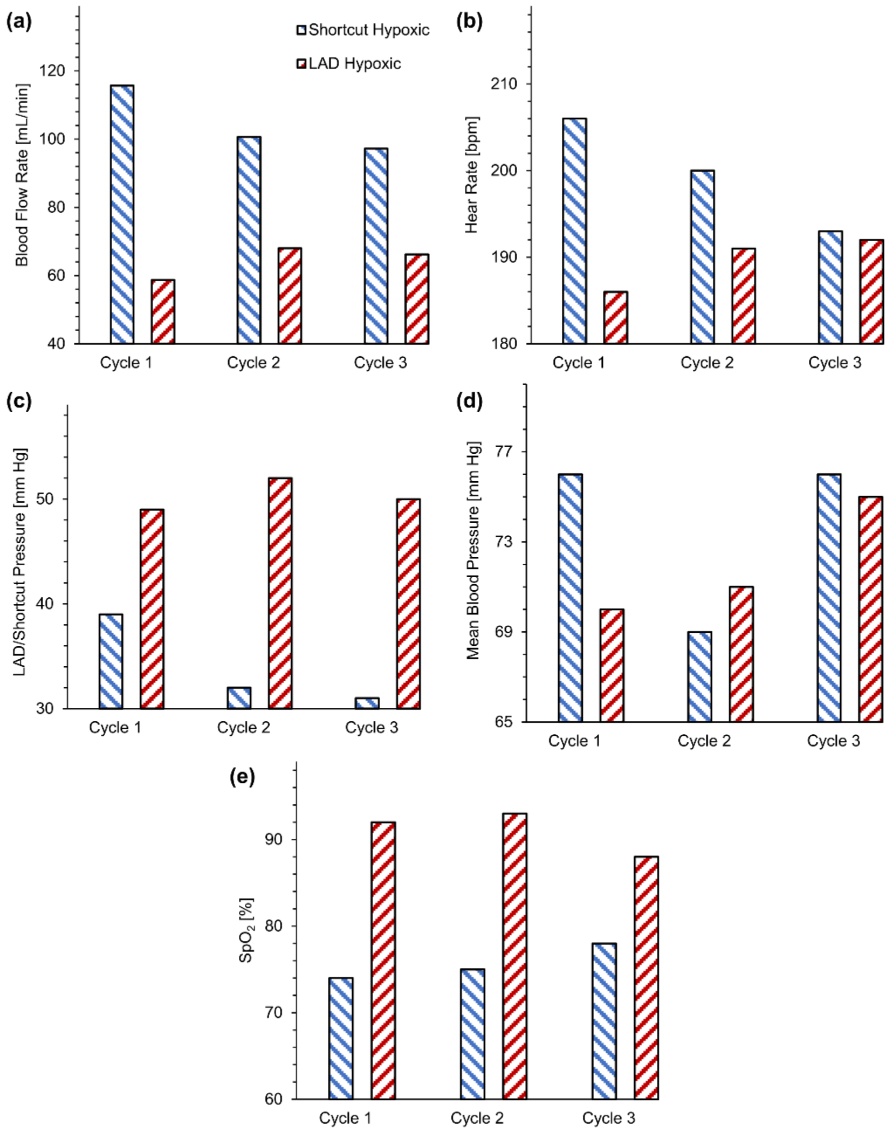


*Figure S10: Effect of extracorporeal bypass on cardiovascular parameters for the LAD with a microchannel height of 130 μm: (a) blood flow rate while the LAD was using oxygen as the sweep gas, (b) heart rate while the LAD was using oxygen as the sweep gas, (c) mean LAD/shortcut pressure while the LAD was using oxygen as the sweep gas, (d) mean systemic arterial blood pressure measured at the femoral artery while the LAD was using oxygen as the sweep gas, and (e) pulse oximeter saturation level.*

### Gas exchange parameters for the LAD with a microchannel height of 130 µm and the piglet

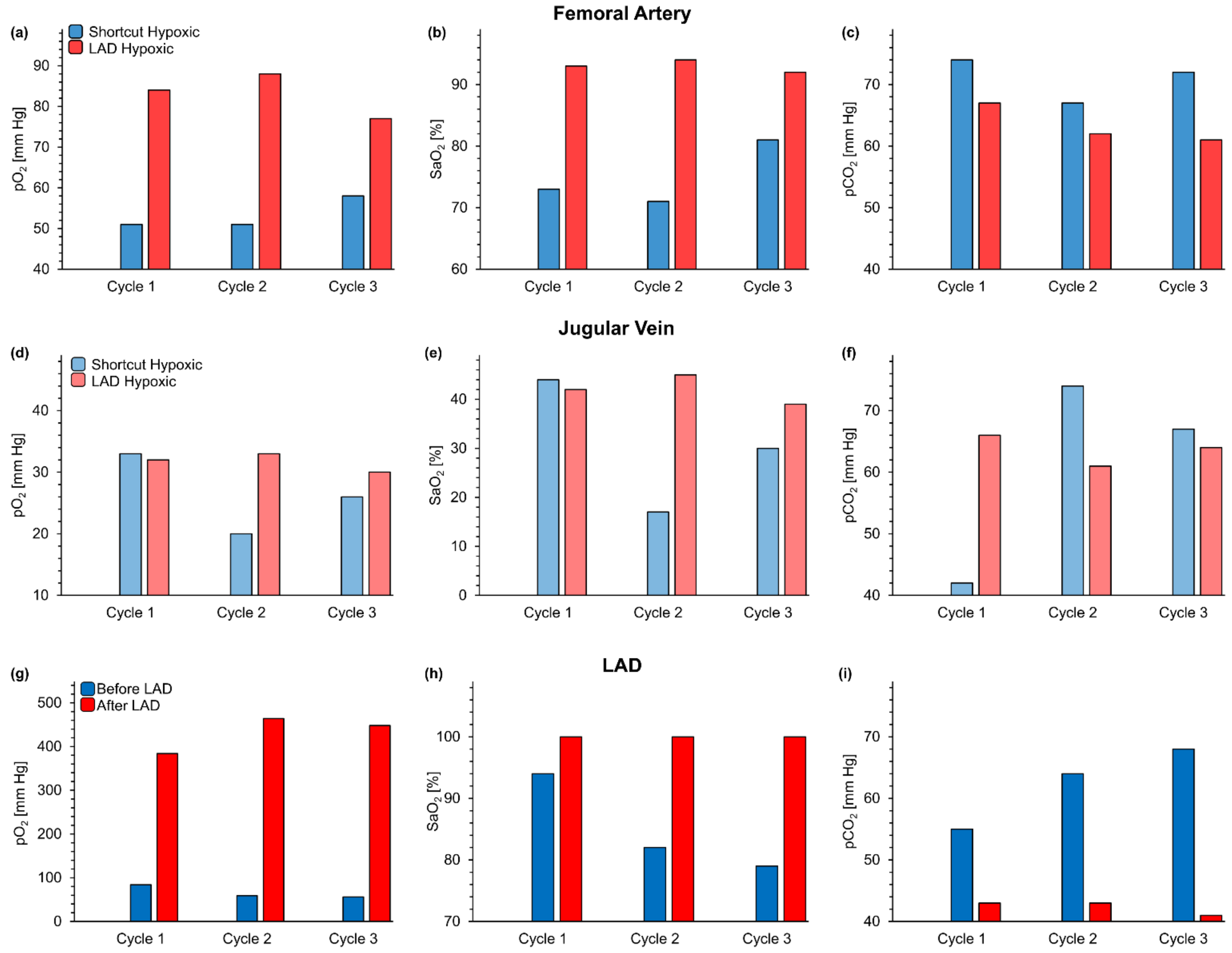


*Figure S11: Gas exchange at femoral artery, jugular vein, and the LAD with a microchannel height of 130 µm when connected in-vivo to a piglet. Measurements were of the blood at the inlet and outlet of the LAD when it was connected to the piglet and pumped by the arterio-venous pressure difference: (a) $pO_2$,(b) Sa $O_2$, (c) $pCO_2$ at femoral artery, (d) $pO_2$, (e) Sa $O_2$, (f) $pCO_2$ at jugular vein, and (g) $pO_2$,(h) Sa $O_2$, (i) $pCO_2$ before and after the LAD. Pure oxygen was used as the sweep gas.*

#### Metabolic analysis by sampling from the femoral artery for the LAD with a microchannel height of 130 µm and the piglet

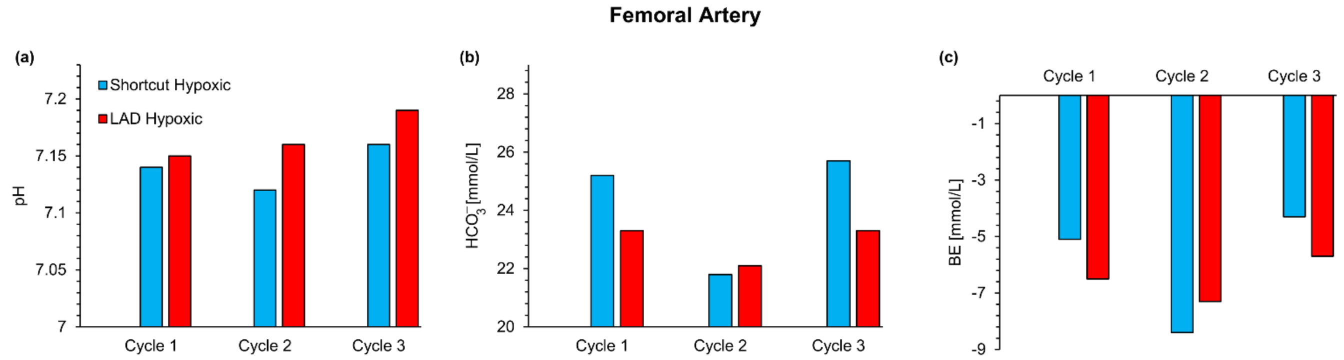


*Figure S12: The metabolic analysis: (a) pH measurement, (b) bicarbonate content, and (c) base excess $pCO_2$ for femoral artery using oxygen as the sweep gas.*

#### Photos of in-vivo experimental setup

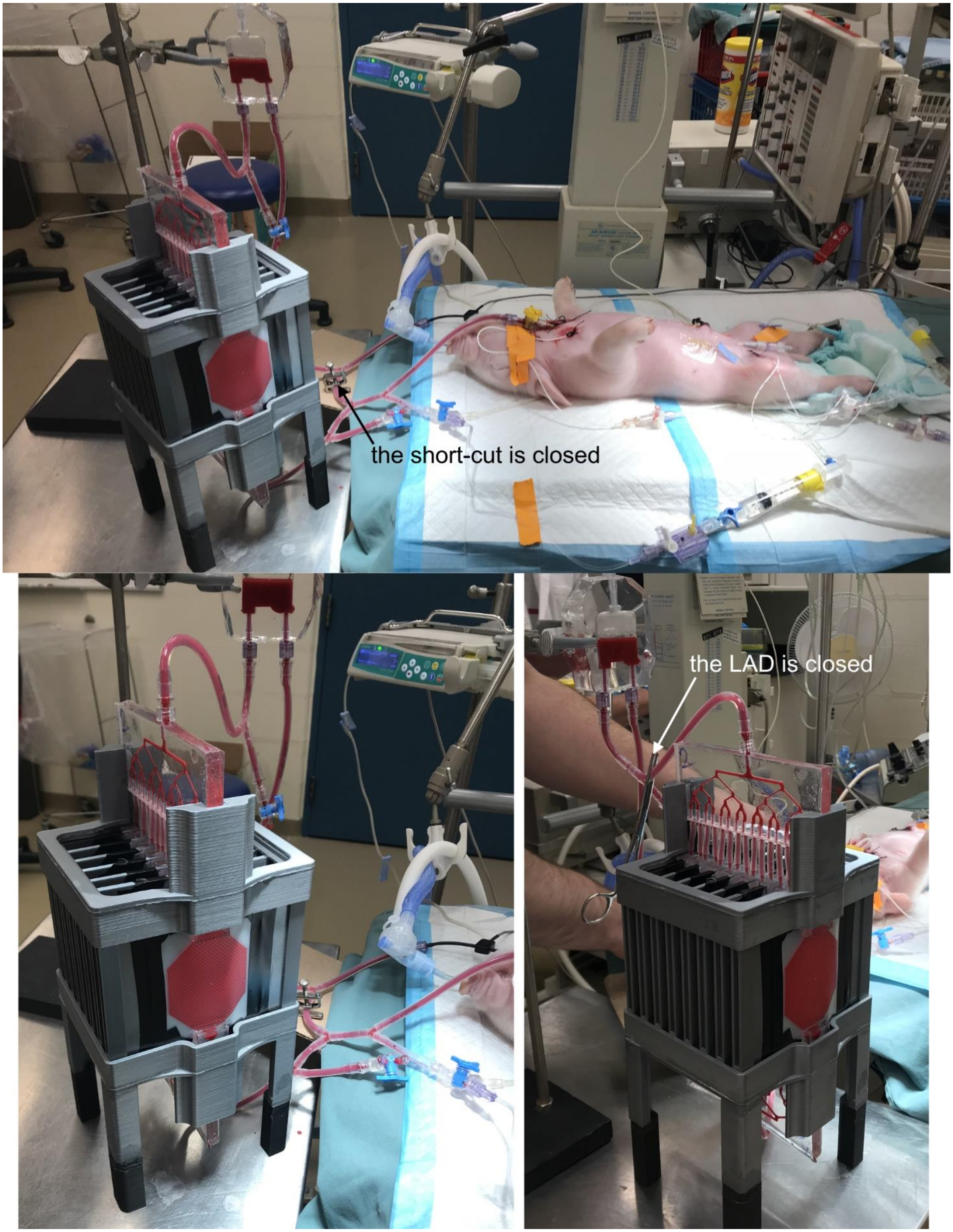


*Figure S13: photos from the animal experiments.*
